# Simultaneous measurement of intrinsic promoter and enhancer potential reveals principles of functional duality and regulatory reciprocity

**DOI:** 10.1101/2025.03.14.643265

**Authors:** Mauricio I. Paramo, Alden King-Yung Leung, Sagar R. Shah, Junke Zhang, Nathaniel D. Tippens, Jin Liang, Li Yao, Yiyang Jin, Xiuqi Pan, Abdullah Ozer, John T. Lis, Haiyuan Yu

## Abstract

Accumulating evidence indicates that both promoters and enhancers are capable of exerting promoter and enhancer functions; however, the relationship between these two activities within individual elements and the determinants of this dual functionality remain poorly understood. We developed a massively parallel dual reporter assay that enables simultaneous assessment of the intrinsic promoter and enhancer potential exerted by the same sequence. Parallel quantification for thousands of elements reveals that canonical human promoters and enhancers can act as both promoters and enhancers under the same contexts, and that promoter activity may be necessary but not sufficient for enhancer function. Perturbations to element transcription factor binding motifs lead to disruptions in both activities, implicating a shared syntax for the two regulatory functions. Combinations of elements with different minimal promoters reveal reciprocal activity modulation, which, together with a strong correlation between promoter and enhancer functions, imply a bidirectional feedback loop to sustain environments of high transcriptional activity. Finally, we validate this reciprocity and correlation *in situ* using CRISPR activation at the human *β*-globin locus. Our approach reveals that the functional convergence between promoters and enhancers arises from a shared regulatory logic and sequence syntax, advancing a unified model for regulatory element biology.

## Main

Cellular processes are precisely controlled through complex gene expression programs that integrate signals from gene-proximal promoters and gene-distal enhancers. Traditionally, promoters and enhancers have been considered separate classes of transcriptional regulatory elements (TREs), often distinguished by their functions, chromatin environments, and genomic locations. Increasing evidence, however, has revealed broad similarities between both the architecture and function of these two regulatory elements, complicating their classification as distinct types^1,2^.

Promoters are classically defined as DNA sequences that drive RNA polymerase II (RNAPII) transcription initiation at gene-proximal transcription start sites (TSS)^3^. The region around the TSS (typically defined as the ±50 base pair (bp) region surrounding the TSS) is known as the core promoter, which contains core promoter motifs, such as the Goldberg-Hogness box (TATA box) and the Initiator (Inr)^4^. These motifs are recognized and bound by general transcription factors (GTFs), such as TBP and TFIIB, that recruit and assemble the RNAPII pre-initiation complex (PIC). RNAPII transcription initiation, promoter-proximal pausing, and pause-release into productive elongation are further facilitated by gene-, cell-, and state-specific transcription factors (TFs) and co-factors that bind at the promoter and at other regulatory sequences such as enhancers to regulate gene transcription^5^.

Conversely, enhancers are defined as DNA sequences that stimulate transcription at a distance, irrespective of their position and orientation with respect to their target gene^6^. The first enhancer discovered was a 72 bp sequence from the SV40 genome that could dramatically increase the transcription of the *β*-globin gene in a distance- and orientation-independent manner^7-9^. Enhancers were soon after discovered in Mammalia^10^ and are now recognized as key elements in the regulation of eukaryotic transcription. Today, enhancers are generally thought of as TF-binding regulatory regions that come into proximity to their target gene promoters via three-dimensional DNA looping^11^.

Numerous findings now point to promoters and enhancers being far more similar than previously thought, spearheaded by the discovery of widespread transcription at enhancer loci^12,13^. Detailed characterization has revealed that when active, both are encompassed within nucleosome-depleted regions (NDRs), bound by sequence-specific TFs that facilitate the assembly of PICs at core promoter regions, and undergo divergent transcription initiation and pausing, with protein-coding gene promoters divergently transcribing mRNAs and upstream antisense RNAs (uaRNAs) in either direction and enhancers divergently transcribing enhancer RNAs (eRNAs) in both^14-19^. These remarkable similarities in chromatin and sequence architecture have led to a proposed unified model for transcriptional regulatory elements^2,14,16^.

Prior attempts at defining promoters and enhancers have largely assumed that their functions are distinct and non-overlapping. However, enhancers possess intrinsic promoter potential^12,13,20^, and numerous studies are now reporting that promoters can exert distal enhancer activity^21-24^. Understanding how these two activities within the same element relate and elucidating their shared determinants is necessary to ensure that current models of gene regulation capture the full complexity of regulatory element functionality.

To date, experiments that measure both the promoter and enhancer potentials of the same set of sequences are lacking. One of the few studies to have employed such a strategy on a large scale was Nguyen et al., which used separate massively parallel reporter assays (MPRAs) to measure promoter and enhancer activities for the same elements^25^. This attempt, however, like all existing MPRAs, inherently decouples the intrinsic promoter and enhancer potential from one another, regulatory potentials that are likely interrelated in their native contexts. One method developed by Mikhaylichenko et al. detected promoter and enhancer activities simultaneously, though was limited to a small test set of elements given the utilization of fluorescent *in situ* hybridization (FISH) as their functional readout^26^.

For these reasons, we developed a Quantitative Unifying Assay for Simultaneously Active Regulatory Regions by sequencing (QUASARR-seq), a massively parallel dual reporter assay that allows for the simultaneous measurement of intrinsic promoter and enhancer potentials exerted by the same element, for thousands of elements in a single experiment. We perform systematic functional comparisons between intrinsic promoter and enhancer functions exerted by canonical promoter and enhancer elements, revealing a degree of entanglement between the two activities, across element types. Our findings suggest that regulatory potential is embedded within element sequences, with impairments in element promoter activity showing compounding effects by also impairing enhancer function. Finally, using combinations of elements with different minimal promoters (minPs), we find that paired elements exert reciprocal regulatory influences to establish promoter and enhancer functions of each element, and validate this phenomenon *in situ* using CRISPR activation (CRISPRa) data at a human enhancer-promoter pair. Our results support the principle that promoters and enhancers are not separate classes of regulatory elements, with their functional convergence underpinned by shared regulatory logic and sequence syntax.

## Results

To investigate the relation and determinants of promoter and enhancer element dual functionality, we sampled a set of TREs in the human immortalized myelogenous leukemia cell line K-562. We previously demonstrated that Precision Run-On and capped nascent RNA sequencing (PRO-cap), a technique that maps TSSs genome-wide by capturing and sequencing the 5′ capped ends of nascent transcripts, provides high predictive performance using 500 bp distance cutoffs from GENCODE annotations, a high-quality, comprehensive reference database of human genes, to distinguish promoter and enhancer elements (see **Methods**)^27,28^. Thus, we selected candidate active promoters defined as GENCODE-proximal PRO-cap TSSs and candidate active enhancers as distal TSSs (**Fig. 1a**). Additionally, we selected a set of candidate inactive promoters defined as proximal PRO-cap untranscribed DNase-seq DNase I hypersensitive sites (DHSs) and candidate inactive enhancers as distal untranscribed DHSs. In total, we tested 2,151 elements, ranging from 65 to 703 bp, with a median length of 254 bp (**Supplementary Fig. 1a**).

**Fig. 1:**
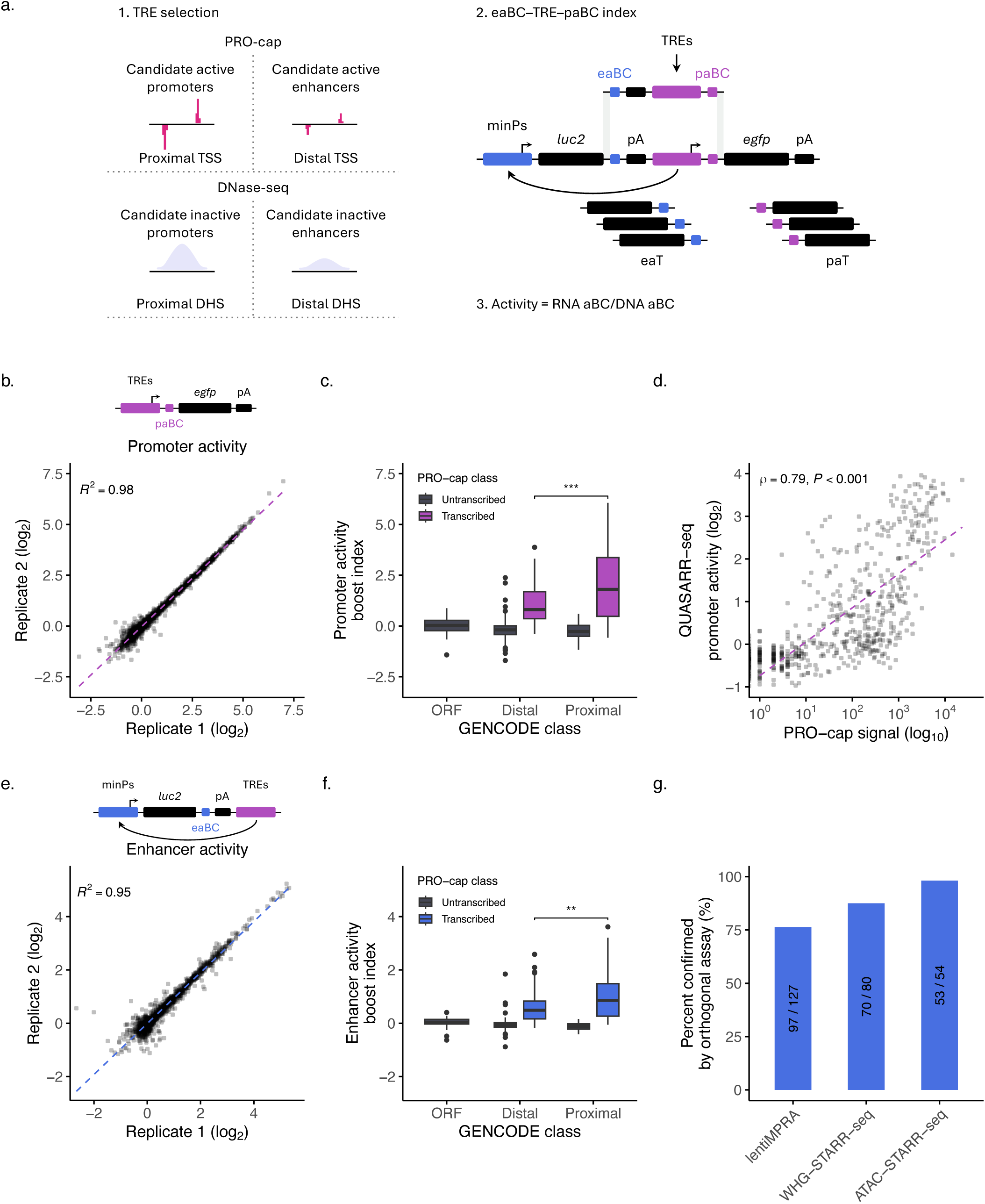
QUASARR-seq simultaneously measures an element’s intrinsic promoter and enhancer activity. **a**, Schematic of the QUASARR-seq workflow. 1, Candidate active promoters were defined as GENCODE-proximal PRO-cap TSSs, active enhancers as distal TSSs, inactive promoters as proximal PRO-cap untranscribed DNase-seq DHSs, and inactive enhancers as distal untranscribed DHSs (see **Methods**). PRO-cap/DNase-seq peak heights reflect typically higher signals at promoters than enhancers. 2, High-complexity eaBC-TRE-paBC indices for accurate and robust measurements are achieved through the generation of eaBC-TRE-paBC amplicons via consecutive isothermal oligo extensions. The QUASARR-seq vector features a minimal promoter (minP; pMYC) that drives the expression of a reporter gene (*luc2*) that contains a degenerate 20 bp enhancer activity barcode (eaBC) located within the 3′ untranslated region (UTR) of an enhancer activity transcript (eaT). Candidate TREs are cloned downstream of this reporter system such that they can drive the expression of a second reporter gene (*egfp*) that contains another degenerate 20 bp promoter activity barcode (paBC) located within the 5′ UTR of a promoter activity transcript (paT). Each reporter system is insulated with a cleavage and polyadenylation site (pA) to avoid confounding the two readouts. 3, QUASARR-seq quantifies activity as the ratio of aBC RNA transcripts to aBC DNA input, normalized using negative controls (see **Methods**). **b**, Correlation of QUASARR-seq promoter activity measurements between replicates. **c**, Promoter activity boost indices of elements parsed by GENCODE and PRO-cap class (*P*-value < 0.001, Student’s t-test). **d**, Correlation between QUASARR-seq promoter activity measurements and PRO-cap signal (Spearman’s *ρ* = 0.79, *P*-value < 0.001). **e**, Correlation of QUASARR-seq enhancer activity measurements between replicates. **f**, Enhancer activity boost indices of elements parsed by GENCODE and PRO-cap class (*P*-value < 0.01). **g**, Percent confirmed of QUASARR-seq active enhancers by lentiMPRA, WHG-STARR-seq, and ATAC-STARR-seq. Calculated based on 50% reciprocal overlap between elements across assays. For box plots, center line represents median, while box limits indicate upper and lower quartiles. Whiskers extend to 1.5 × interquartile range, and points beyond whiskers denote outliers. Significance between groups was assessed using Student’s t-test, with significance levels indicated by asterisks (* *P*-value < 0.05, ** *P*-value < 0.01, *** *P*-value < 0.001).

Hereafter, for clarity, we use the terms ‘promoter activity’ to refer to the candidate TREs ability to drive transcription locally (i.e., initiation at the TRE) and ‘enhancer activity’ to refer to the candidate TREs ability to drive transcription from a distance (i.e., initiation at the minP).

### QUASARR-seq simultaneously measures an element’s intrinsic promoter and enhancer activity

To systematically evaluate the functional duality of transcriptional regulatory elements, we developed a Quantitative Unifying Assay for Simultaneously Active Regulatory Regions by sequencing (QUASARR-seq), a massively parallel dual reporter assay that enables the simultaneous measurement of intrinsic promoter and enhancer potentials exerted by the same regulatory element molecule (**Fig. 1a**). QUASARR-seq adopts the principles of classical reporter assays where candidate regulatory sequences are cloned into expression vectors but incorporates two functional reporter systems each used to independently quantify promoter and enhancer activity for the same sequence (see **Methods**). Enhancer activity measurements are obtained by the reporter system positioned upstream of the TRE that contains a minP that drives the expression of enhancer activity barcodes (eaBCs) located within the 3′ untranslated region (UTR) of an enhancer activity transcript (eaT). Similarly, promoter activity measurements are obtained by the reporter system positioned downstream of the TRE where the TRE itself drives the expression of promoter activity barcodes (paBCs) located within the 5′ UTR of a promoter activity transcript (paT). As in all MPRAs, QUASARR-seq quantifies activity as the ratio of counts of aBC RNA transcripts to aBC DNA input vectors, from which boost indices are calculated by basal expression level normalization using negative controls (see **Methods**).

QUASARR-seq promoter activity measurements [log_2_(RNA paBCs/DNA paBCs)] across replicates demonstrated a high degree of reproducibility (*R^2^* = 0.98; **Fig. 1b**). As expected, promoter activity varied significantly across GENCODE classes, with PRO-cap transcribed proximal elements (i.e., active promoters) exhibiting the highest activity, followed by transcribed distal elements (i.e., active enhancers), and with untranscribed elements and negative control open reading frames (ORFs) exhibiting little or no activity, respectively (*P*-value < 0.001, Student’s t-test; **Fig. 1c**). Interestingly, many of the proximal elements with the highest promoter activities were found to correspond to the promoters of long non-coding RNAs (lncRNAs) and small nuclear RNAs (snRNAs). The top three included: 1. the divergent promoter for LINC01138 (lncRNA)/RNVU1-21 (snRNA) and ENSG00000224481 (uncharacterized lncRNA), 2. the promoter for RNVU1-2A (snRNA), and 3. the divergent promoter for SNHG1 (lncRNA) and SLC3A2 (protein coding).

Promoter activity measurements showed a good correlation with native genomic PRO-cap (Spearman’s *ρ* = 0.79, *P*-value < 0.001; **Fig. 1d**) and PRO-seq signals analyzed under a variety of criteria (**Supplementary Fig. 2**; see **Methods**), suggesting that QUASARR-seq captures a significant, though incomplete reflection of transcription initiation and elongation patterns observed in endogenous contexts, similar to other reports using transient systems^22,25^. We hypothesize that these differences may be due to the effects of enhancers and other regulatory elements acting on the TREs in the native loci while QUASARR-seq is capturing their intrinsic promoter activity. We observed a similar correlation with Survey of Regulatory Elements^29^ (SuRE; Spearman’s *ρ* = 0.52, *P*-value < 0.001; **Supplementary Fig. 3a**) to those from PRO-seq (**Supplementary Fig. 2**), despite the low 10% reciprocal overlap between elements tested in both assays.

QUASARR-seq enhancer activity measurements [log_2_(RNA eaBCs/DNA eaBCs)] across replicates also demonstrated a high degree of reproducibility (*R^2^* = 0.95; **Fig. 1e**). Enhancer activity too varied across GENCODE classes, albeit to a lesser degree, with proximal elements exhibiting a dynamic range closer to that of distal elements (*P*-value < 0.01; **Fig. 1f**). Again, many of the proximal elements displaying the highest enhancer activities corresponded to lncRNA promoters. The top three included: 1. the promoter for ENSG00000266401 (uncharacterized lncRNA), 2. the divergent promoter for ENSG00000287126 (uncharacterized lncRNA) and RPS7 (protein coding), and 3. the divergent promoter for SNHG1 (lncRNA) and SLC3A2 (protein coding). These results agree with extensive reports finding high levels of enhancer activity mediated by the promoters of lncRNAs^24,30,31^.

QUASARR-seq active enhancer calls were reliably corroborated by orthogonal assays, with ATAC-STARR-seq (98.1%), WHG-STARR-seq (87.5%), and lentiMPRA (76.4%) all showing reciprocal active enhancer call rates greater than 75% (**Fig. 1g**). Similarly, QUASARR-seq reliably captured active enhancer calls made by orthogonal assays (**Supplementary Fig. 3b**). In all, QUASARR-seq enables the simultaneous quantification of both promoter and enhancer activities from the same element, uniquely positioning us to address long-standing questions about the dual functionality of transcriptional regulatory elements.

To gain insight into the functional balance between activities, we calculated the balance index of promoter-to-enhancer activity boost indices (see **Methods**). This balance index was used to determine if an element predominantly acted as a promoter, an enhancer, or had a balanced dual function. Since promoter activity had a wider dynamic range than enhancer activity (**Supplementary Fig. 4a**), we Z-score normalized the two measurements prior to calculating the balance index. We defined "Promoter-dominant" as elements with a balance index > 1 (stronger promoter activity relative to enhancer activity), "Enhancer-dominant" as elements with a balance index < -1 (stronger enhancer activity relative to promoter activity), and "Balanced" as elements with a balance index falling between -1 and 1 (similar levels of promoter and enhancer activity, that is, a balanced function). We used these standard cutoffs as they corresponded to one standard deviation from the mean, highlighting elements with relatively strong activity dominance.

The distribution of balance indices differed significantly across GENCODE classes (*P*-value < 0.001, Student’s t-test; **Supplementary Fig. 4b**), with the number of elements in different balance index categories varying between proximal and distal elements (**Supplementary Fig. 4c**). Proximal elements were mostly balanced (61.4%), but with a considerable number also displaying promoter-dominant behavior (34.9%). Interestingly, distal elements were predominantly balanced (86%), with very few enhancer-dominant (8.3%), indicating that promoter potential may be required for enhancer function.

To investigate the potential drivers of functional bias in activity balance, we calculated the GC content (see **Methods**) of elements to compare between balance index categories. For both proximal and distal TREs, promoter-dominant elements were significantly more GC-rich than their balanced (proximal and distal, *P*-value < 0.001, Student’s t-test with Bonferroni correction) and enhancer-dominant (proximal, *P*-value < 0.01; distal, *P*-value < 0.001) counterparts (**Supplementary Fig. 4d-e**), suggesting that activity preference is, at least partially, sequence encoded. These results are consistent with previous observations that CpG-rich sequences show elevated promoter activity and promoter-to-enhancer ratios^25^, and further indicate that GC content drives promoter-dominant functional bias across element classes, revealing a unifying sequence syntax beyond classical promoter or enhancer demarcations.

Next, to investigate the influence of core promoter architecture on dual regulatory activity, we calculated motif match scores (see **Methods**) for elements within specified windows relative to their GRO-cap-detected TSSs in K-562, including TATA (-40 to -20 bp), Inr (-2 bp), MTE (+5 to +35 bp), and DPE (+5 to +35 bp). To assess whether specific core promoter motifs contribute to establishing either activity bias or element class, we compared motif match scores with balance index categories across GENCODE classes. As expected, the distributions for proximal and distal TREs showed similar relationships between core promoter motif match scores and balance index categories (**Supplementary Fig. 4f**), consistent with previous evidence of a unified sequence architecture for promoters and enhancers^14^. Still, some differences emerged, primarily across balance index categories. In proximal TREs, promoter-dominant elements had significantly higher Inr and MTE scores than balanced elements (*P*-value < 0.05, Wilcoxon rank-sum test with Bonferroni correction). In distal TREs, promoter-dominant elements also had higher Inr scores than balanced (*P*-value < 0.01) in addition to enhancer-dominant (*P*-value < 0.001) elements. By contrast, promoter-dominant elements had lower TATA scores than balanced (*P*-value < 0.001) and enhancer-dominant (*P*-value < 0.01) elements, whereas DPE was enriched in balanced relative to enhancer-dominant elements (*P*-value < 0.05).

Together, these data suggest that GC content and Inr are the most informative sequence predictors of promoter-biased activity across proximal and distal TREs. In contrast, TATA is a weak predictor of promoter dominance, likely due to strong TATA-containing elements making up only ∼10% of TREs. MTE and DPE showed modest, context-dependent contributions, influencing activity bias only in specific TRE classes.

### Promoter and enhancer activities are positively correlated

Both positive and negative relationships between promoter and enhancer activities have been proposed^2,12^. According to the positive relationship hypothesis, the strength of an element’s promoter activity is directly related to its enhancer activity. In contrast, in the negative relationship hypothesis, the individual promoter activity strengths of interacting elements determine their respective enhancer activities following an inverse relationship, likely established by competition between the elements to determine their primary activity in a context-dependent manner.

Prior attempts at elucidating this relationship used decoupled promoter and enhancer activity measurements^25^, making the negative relationship hypothesis untestable as they could not capture the dynamic interplay between promoter and enhancer functions within the same element. Since the negative relationship hypothesis posits mutual exclusivity in promoter and enhancer activities, assessing their interrelation requires simultaneous measurement of both functions. To address this limitation, we utilized activity measurements obtained by QUASARR-seq, as it uniquely allows us to perform an agnostic comparison between these two competing models. QUASARR-seq measured promoter and enhancer activities across TREs were substantially positively correlated (Spearman’s *ρ* = 0.89, *P*-value < 0.001; **Fig. 2a**), more so than prior reports that yielded similar results^25^. These data support a model where the ability to drive transcription locally (i.e., promoter activity) may serve as a proxy for the ability to drive transcription from a distance (i.e., enhancer activity).

**Fig. 2:**
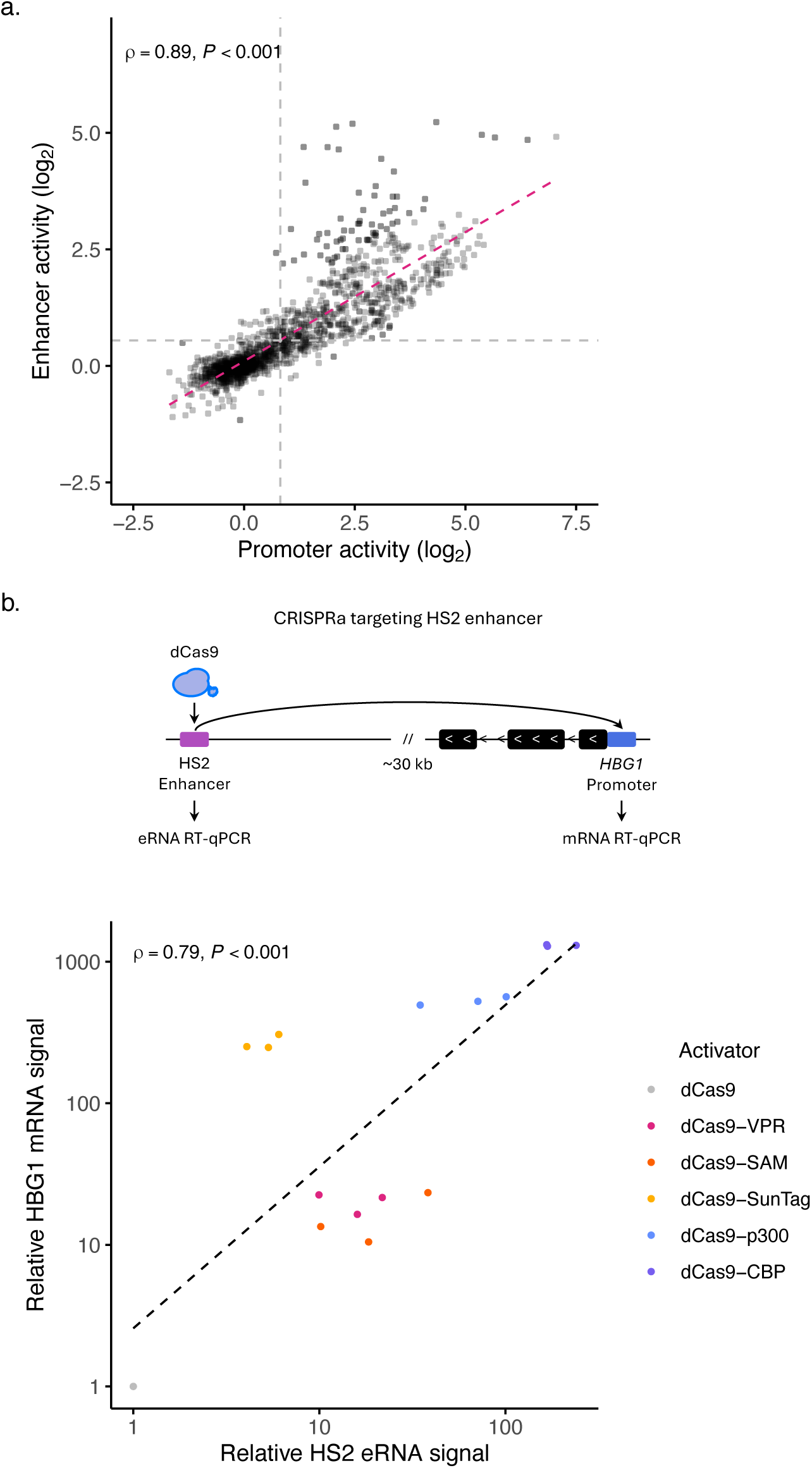
Promoter and enhancer activities are positively correlated. **a**, Correlation between QUASARR-seq measured promoter and enhancer activities (Spearman’s *ρ* = 0.89, *P*-value < 0.001). **b**, Correlation between RT-qPCR measured HS2 eRNA signal and its target gene promoter HBG1 mRNA signal (Spearman’s *ρ* = 0.79, *P*-value < 0.001) when HS2 enhancer is targeted with different dCas9-based transcriptional activators, including dCas9-VPR, dCas9-SAM, dCas9-SunTag, dCas9-p300, dCas9-CBP, and dCas9 as a negative control. Points represent signal relative to dCas9. CRISPRa data from Wang et al.^32^

Despite each reporter system being insulated with a cleavage and polyadenylation site (pA) to avoid confounding the two activity readouts (**Fig. 1a**; see **Methods**), we asked whether the high correlation between promoter and enhancer activities that we observed may be due to measuring the same function, perhaps from pA read-through transcription. To examine this, we performed RT-qPCR targeting *luc2* (upstream of the first pA) and *egfp* (downstream of the first pA) in both TRE-containing and TRE-lacking empty vector constructs (see **Methods**). Across experiments, we found that read-through transcription accounted for 16.47% of *egfp* expression, indicating that a relatively small fraction of the promoter activity signal measured reflects transcriptional leakage past the first pA (**Supplementary Fig. 5a**). These data suggest that QUASARR-seq promoter and enhancer activity measurements are primarily capturing related, but distinct regulatory functions of the same elements.

We next sought to assess whether the positive relationship between promoter and enhancer functions observed in QUASARR-seq could be validated in an endogenous genomic context. To this end, we re-analyzed a published CRISPRa dataset in which the enhancer HS2 was targeted with different dCas9-based transcriptional activators in HEK293T, including dCas9-VPR, dCas9-SAM, dCas9-SunTag, dCas9-p300, dCas9-CBP, and dCas9 as a negative control^32^. In this *in situ* context, the enhancer HS2 is located ∼30 kb downstream of, and convergent with, its target gene promoter *HBG1*. Given that activators exhibit distinct activation strengths, we examined whether varying levels of enhancer transcription activation were accompanied by corresponding changes in target promoter transcription. Targeting of HS2 with the different activators showed a strong positive correlation between HS2 eRNA activation (i.e., HS2 promoter activity) and HBG1 mRNA activation (i.e., HS2 enhancer activity, Spearman’s *ρ* = 0.79, *P*-value < 0.001; **Fig. 2b**). Together with our findings, these data indicate that the positive correlation between promoter and enhancer activities holds across endogenous genomic contexts and distances.

### Promoter activity is necessary, but not sufficient for enhancer function

While increasing evidence suggests that transcriptional regulatory elements can exert both promoter and enhancer activities, it is still unclear whether these functions can co-occur or are mutually exclusive under the same regulatory contexts. Since QUASARR-seq measures both activities exerted by the same sequence simultaneously, we asked whether elements with measured promoter activity were also capable of being assayed for enhancer function.

To investigate the concurrence of activities, we used a uniform processing pipeline (see **Methods**) to obtain QUASARR-seq-derived promoter and enhancer calls to make comparisons between activities. For this analysis, we used only data where we had paired promoter and enhancer activity measurements for the same element from the same experiment. Of the QUASARR-seq-called active promoters, we found that 69% (180/261) were also called active enhancers (**Fig. 3a**). More strikingly, of the QUASARR-seq-called active enhancers, 96.3% (180/187) were also called active promoters. These results suggest that promoter and enhancer activities can co-occur within the same regulatory context and that promoter activity may be necessary, but not sufficient for enhancer function.

**Fig. 3:**
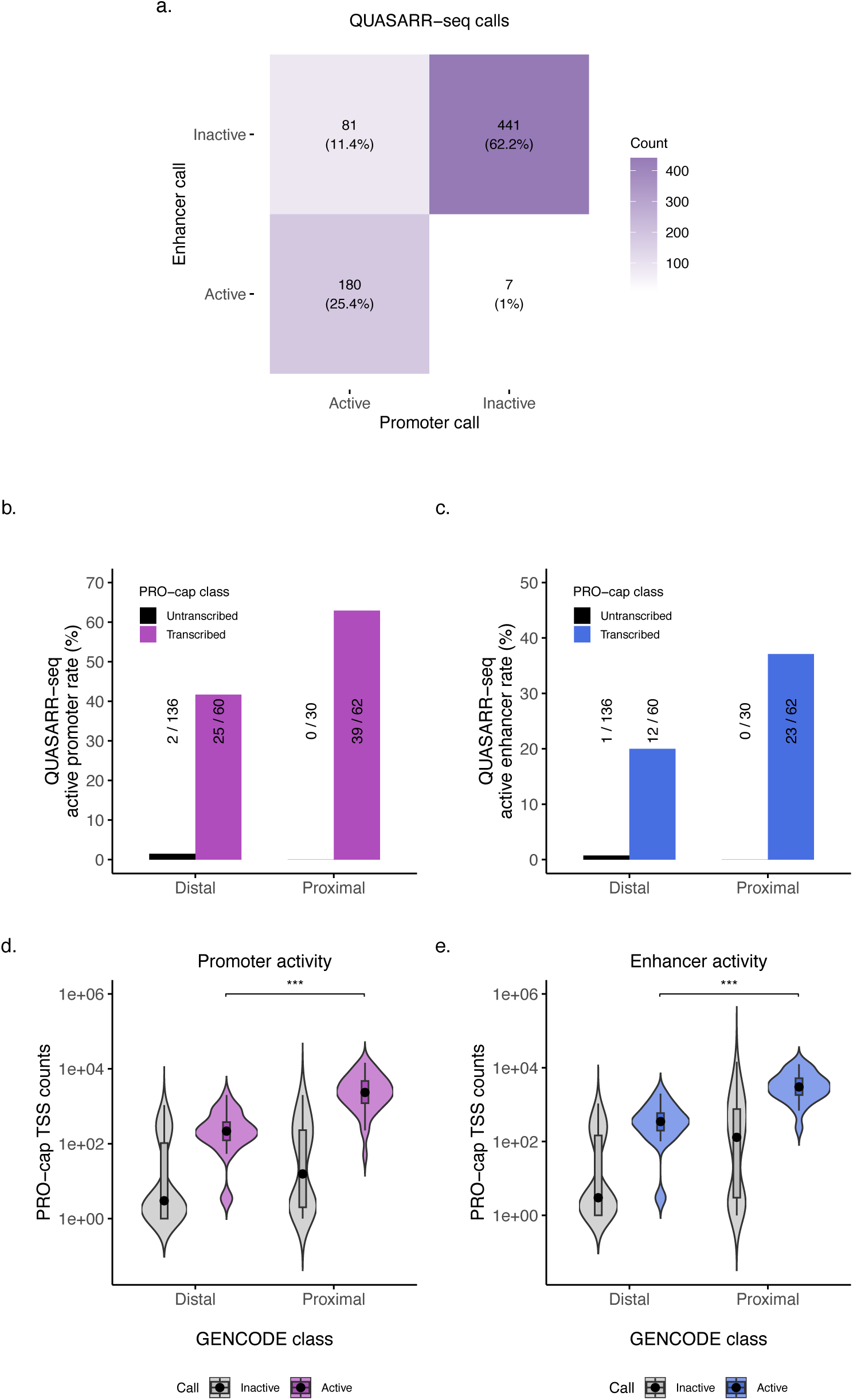
Promoter activity is necessary, but insufficient for enhancer function. **a**, Heatmap showing overlap between QUASARR-seq promoter and enhancer active element calls. Only elements with paired promoter and enhancer activity measurements obtained from the same experiment are included in this analysis. **b-c**, QUASARR-seq active promoter (b) and enhancer (c) rates by GENCODE and PRO-cap class. **d-e**, Element PRO-cap TSS counts by GENCODE class and QUASARR-seq active promoter (d) and enhancer (e) call (*P*-value < 0.001, Student’s t-test).

### Promoters and enhancers can perform both promoter and enhancer functions

We next asked whether QUASARR-seq active calls would stratify elements based on their canonical classifications. While almost all QUASARR-seq active promoters were PRO-cap transcribed elements, not all PRO-cap transcribed elements were deemed QUASARR-seq active promoters (**Fig. 3b**). This suggests that QUASARR-seq exhibits a high degree of specificity, with some reduction in sensitivity, which we hypothesize may be due to capturing the element’s promoter activity without the effects of other regulatory elements acting on them at their native loci. Similarly, essentially all QUASARR-seq active enhancers were PRO-cap transcribed (**Fig. 3c**), in line with our previous report that showed that transcription serves as a robust predictor for active enhancer elements^27^.

Both canonical promoters and enhancers could perform both promoter and enhancer functions. Still, active promoter calls varied across GENCODE classes, with proximal elements displaying the highest active rate (62.9%; **Fig. 3b**), followed by distal elements (41.7%). We observed a similar trend with active enhancer calls, with proximal elements exhibiting the highest active rate (37.1%; **Fig. 3c**), followed by distal elements (20%).

To explore potential drivers of differences in promoter and enhancer activity rates across GENCODE classes, we calculated the total PRO-cap read counts within the boundaries of tested elements. We hypothesized that the disparity in activity rates might reflect the inherent ability of proximal elements to initiate transcription at higher frequencies than distal elements. Supporting this hypothesis, we found that proximal elements generally exhibited higher endogenous initiation frequencies compared to distal elements (*P*-value < 0.001, Student’s t-test; **Fig. 3d-e**). Moreover, all active promoter and enhancer elements trended toward the upper bound in TSS counts.

In all, these data strengthen previous associations between transcription and enhancer activity. We find compelling evidence that essentially all active elements, independent of traditional classifications, require some intrinsic ability to initiate transcription. This suggests a functional entanglement in both activities, present across element types.

### Downstream sequence features do not confer element activity or type

Despite overwhelming similarity in chromatin and sequence architecture, the RNA transcripts produced by promoters and enhancers exhibit striking differences in their properties^20,33-35^ (**Fig. 4a**). mRNAs are typically long, spliced, and polyadenylated, whereas eRNAs are short, unspliced, and non-polyadenylated. Extensive evidence suggests that these differences in RNA classes are largely determined by the sequences present at the 5′ ends of transcripts (i.e., immediately downstream of the elements). Despite being non-polyadenylated, utilization of early poly(A) site sequences (e.g., AT/ATAA) is a discerning feature of eRNA transcription, which contributes to their short length and reduced half-life due to rapid degradation by the exosome complex^36^. In contrast, mRNAs are enriched with U1 small nuclear ribonucleoprotein (snRNP) recognized 5′ splice sites and are depleted of early poly(A) sites, where early U1 binding appears to impede early termination^37-40^.

**Fig. 4:**
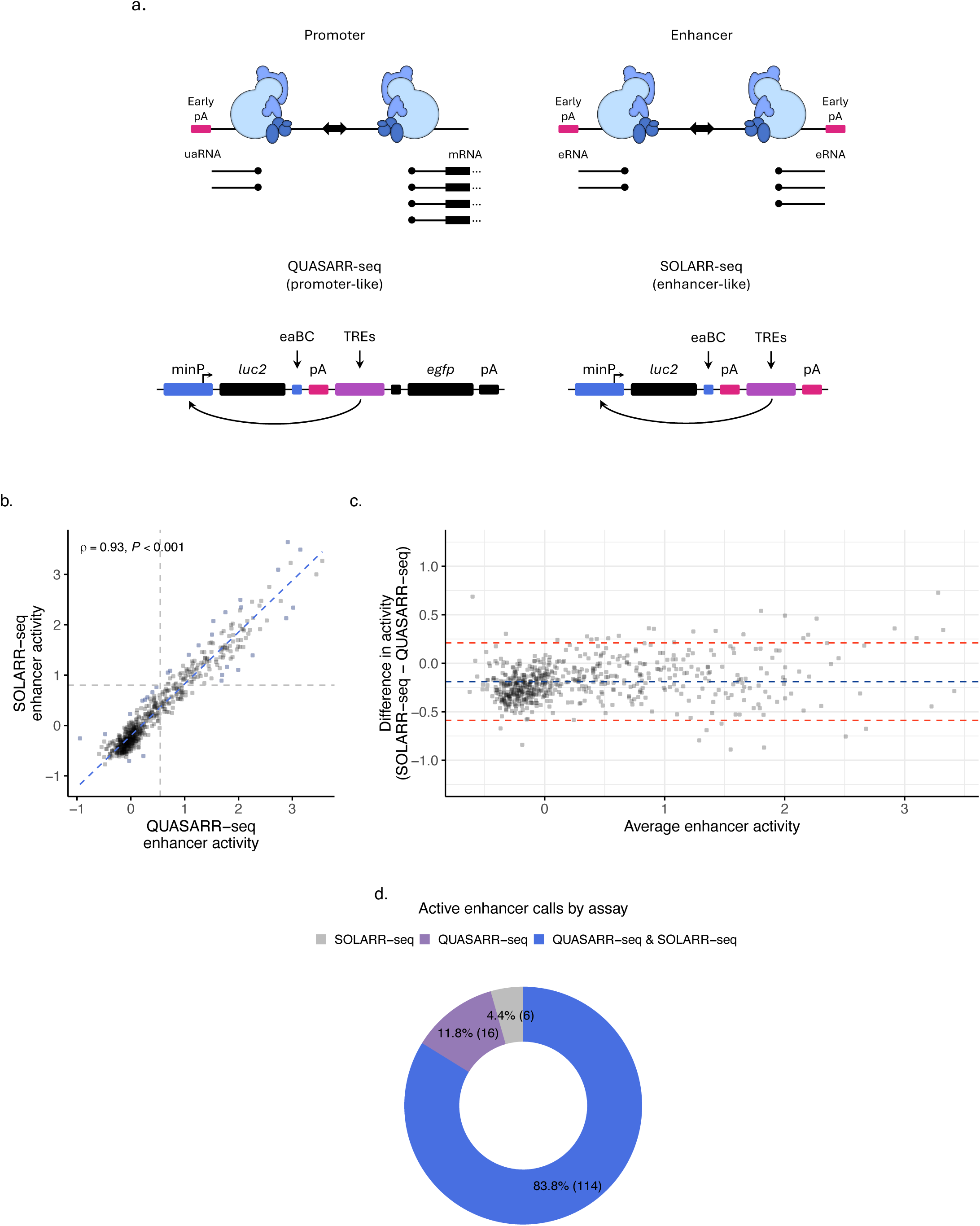
Downstream sequence features do not confer element activity or type. **a**, Schematic showcasing differences in downstream sequence features between canonical promoters and enhancers. QUASARR-seq mimics downstream features to those of promoters while SOLARR-seq mimics downstream features to those of enhancers. **b**, Correlation between QUASARR-seq and SOLARR-seq measured enhancer activities (Spearman’s *ρ* = 0.93, *P*-value < 0.001). **c**, Bland-Altman plot of agreement showing differences in activity measurements between SOLARR-seq and QUASARR-seq. **d**, Active enhancer calls by QUASARR-seq and SOLARR-seq.

It has been hypothesized that the early termination signals employed during eRNA transcription might contribute to enhancer function by promoting RNAPII turnover, which, presumably, could facilitate the transfer of transcriptional machinery from the enhancer to its associated promoter. In contrast, the absence of early termination signals and the presence of 5′ splice sites in mRNAs may inhibit transcriptional termination, thereby supporting efficient local elongation.

To test the hypothesis that downstream sequence features mediate *enhancerization* of regulatory elements, we compared enhancer activity measurements for the same set of TREs using two assays configured to simulate promoter and enhancer downstream regions. Since QUASARR-seq is designed to measure the intrinsic promoter activity of TREs, its paT is modeled to mimic attributes of mRNAs (**Fig. 4a**). In contrast, we developed Surveying OLigos for Active Regulatory Regions and sequencing (SOLARR-seq), a massively parallel reporter assay identical in sequence to QUASARR-seq except for the absence of the paBC and *egfp* components used for promoter activity measurements. Notably, SOLARR-seq incorporates early pAs immediately downstream of the TRE cloning cassette, mimicking attributes of enhancer downstream regions. We hypothesized that if early termination stimulated enhancer activity, a significant difference in activity would be observed between the two assays’ enhancer activity measurements.

SOLARR-seq enhancer activity measurements between replicates demonstrated a high degree of reproducibility (*R^2^* = 0.92; **Supplementary Fig. 6a**). Interestingly, a strong positive correlation was observed between QUASARR-seq and SOLARR-seq enhancer activity measurements (Spearman’s *ρ* = 0.93; **Fig. 4b**), indicating that despite substantial differences in downstream features, the two assays provided comparable enhancer activity readouts. To evaluate potential systematic biases, including mean differences and outliers, we performed a Bland-Altman analysis^41^. This revealed consistent agreement between the two assays across activity ranges, with most differences falling within the limits of agreement (**Fig. 4c**). Using a uniform processing pipeline (see **Methods**), to obtain active enhancer calls for both QUASARR-seq and SOLARR-seq, we found a substantial overlap of active enhancers identified by both assays (83.8%; **Fig. 4d**), and that both proximal and distal elements were capable of exerting this enhancer function (**Supplementary Fig. 6b**).

Altogether, these results indicate that the striking differences in downstream sequence features between promoters and enhancers do not appear to influence regulatory activity or play a role in establishing element type, suggesting that regulatory potential is likely encoded within element sequences.

### Likely pathogenic variants that impair promoter activity have compounding effects by also impairing enhancer activity

To further investigate the determinants of promoter and enhancer dual functionality, we sought to test the hypothesis that regulatory potential is embedded within element sequences. In previous work, we demonstrated that deletion of TRE core promoter regions (defined as −35 to +60 bp from the TSS) significantly impairs, and often completely ablates enhancer function^27^. Complete deletion of these core promoter regions, which disrupts enhancer function, would almost certainly also disrupt promoter function. Thus, to examine whether impairment of an element’s promoter activity leads to corresponding impairment of its enhancer activity with greater precision, we first focused on testing the impact of variants with strong clinical/functional evidence for disrupting promoter activity.

The variant c.-192A>G (ClinVar ID: 243006; gnomAD: 0.007%) is a nucleotide substitution 192 bp upstream of the ATG translational start site of the *APC* promoter 1B region^42^. This variant has been observed in individuals with clinical features of gastric adenocarcinoma and proximal polyposis of the stomach. Functional studies have demonstrated a damaging effect via binding disruptions of the TF YY1 and impaired activity of the *APC* promoter in gastric and colorectal cancer cell lines. Promoter impairment is further supported by an allelic imbalance in patient blood cells that suggests decreased allele-specific expression *in vivo*. Based on these findings, this variant has been classified as ‘Likely Pathogenic’.

In addition to c.-192A>G, hereafter referred to as A4G, we generated three additional variants (A1C, T5C, and G6A), each targeting a different base of the same YY1 motif (AAG[G/A][T/G]GGC) within the *APC* promoter (**Fig. 5a**). QUASARR-seq measurements of wild-type and mutant alleles of the element corroborated reported promoter activity impairment. More strikingly, simultaneous measurement of enhancer activity revealed a corresponding impairment on enhancer function. These results suggest that disruptions in an element’s promoter activity by likely pathogenic variants can have compounding effects by also disrupting its enhancer function. Dual impairment of regulatory activity could have broader implications in the way that damaging effects of disease-associated variants are identified or potentially treated.

**Fig. 5:**
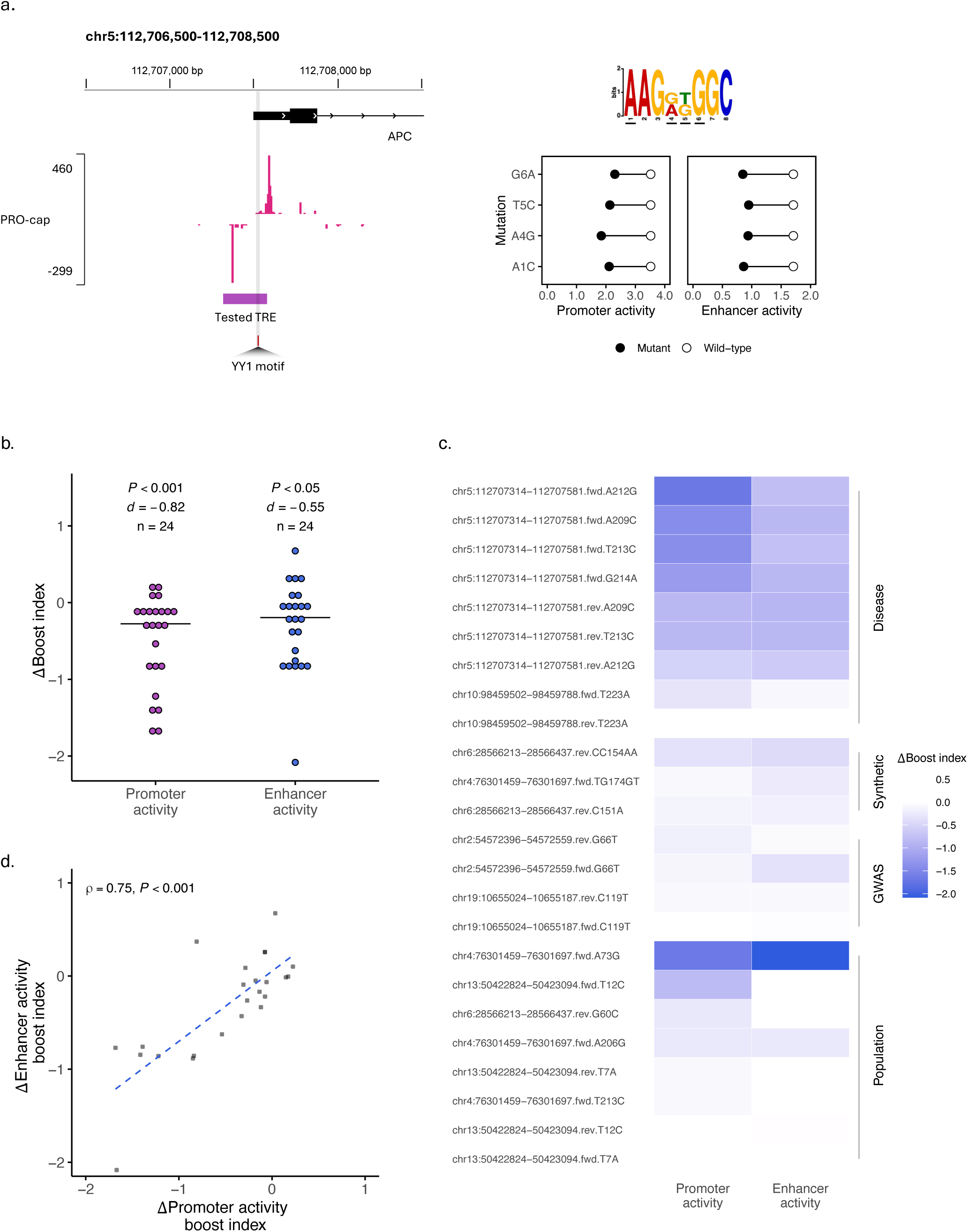
Promoter and enhancer effects are correlated in variants disrupting dual function. **a**, Left, Genome browser of the APC promoter locus that contains likely pathogenic variants. Top right, PWM logo of YY1 motif found in tested TRE. Bottom right. Lollipop plot showing wild-type (white circle) and mutant (black circle) allele promoter (left plot) and enhancer (right plot) activity measurements. **b**, Change (Δ) in promoter and enhancer activity boost indices between mutant and wild-type alleles. Dots represent individual elements (n = 24), horizontal bars indicate median Δ boost indices. Significance assessed using One-sample t-test and effect size as Cohen’s *d*. **c**, Heatmap of Δ in promoter and enhancer activity boost indices for the disease-associated, synthetic, genome-wide association study (GWAS), and population variants tested. **d**, Correlation of the Δ between promoter and enhancer activity boost indices (Spearman’s *ρ* = 0.75, *P*-value < 0.001).

### Promoter and enhancer effects are correlated in variants disrupting dual function

To generalize these results, we expanded the set of elements to assess the effects of additional variants on dual regulatory function. We generated mutant elements containing disease-associated (n = 9), synthetic (n = 3), genome-wide association study (GWAS; n = 4), and population (n = 8) variants. In this analysis, we included only elements where we obtained both the promoter and enhancer activity of the wild-type and mutant alleles.

To assess the relation between impairments in dual regulatory function, we calculated the change (Δ) in promoter and enhancer activity boost indices by subtracting the wild-type element boost index from the mutant element boost index (see **Methods**). Here, a low Δ boost index indicated a large mutant disrupting effect. Across the population, variants significantly reduced QUASARR-seq promoter activity measurements (*P*-value < 0.001, One-sample t-test, Cohen’s *d* = -0.82, large; **Fig. 5b**), although the effect was variable for individual elements. Moreover, we observed a similar trend with enhancer activity measurements, though with a lesser effect size compared to promoter activity (*P*-value < 0.05, *d* = -0.55, medium).

We next asked which mutation classes inflicted the largest disrupting effects on regulatory element activity. As expected, disease-associated variants generally imparted the lowest Δ in dual regulatory function (**Fig. 5c**), with a few population variants unexpectedly showing some high degree of disruption effects. We observed a trend between the Δ promoter and enhancer boost indices and found that both were substantially positively correlated (Spearman’s *ρ* = 0.75, *P*-value < 0.001; **Fig. 5d**). These results indicate that variants that disrupt an element’s promoter activity generally also disrupt its enhancer activity, suggesting a shared syntax for the two regulatory functions.

### Paired elements exert reciprocal regulatory influences to establish activities of both elements

Recent studies propose a model in which an element’s enhancer activity is established by the quantitative tuning of the intrinsic promoter activity of its associated promoter, its own intrinsic enhancer activity, and the three-dimensional contacts between the two, with TRE compatibility appearing to play an additional, though limited role^43^. It should follow then that an element’s promoter-enhancer dual functionality can be shaped by its interactions with other regulatory elements, including by its associated minimal promoter.

To investigate the effects that different minimal promoters have on an element’s promoter and enhancer activities, we performed QUASARR-seq by pairing TREs with different minPs. For these experiments, we included the promoters of a housekeeping gene; *GAPDH*, and a developmental gene; *APOBEC3F*, in addition to the promoter of *MYC* used thus far in this study. To distinguish between libraries, we incorporated a unique three-bp barcode specific to each minP library positioned directly downstream of the paBCs, allowing us to identify the origin of each activity score and attribute it to its associated minP (see **Methods**). QUASARR-seq promoter activity measurements between replicates across minP libraries were highly reproducible (**Supplementary Fig. 7a-c**), with some reduction for enhancer activity measurements relative to promoter activity (**Supplementary Fig. 7d-f**).

Next, we calculated the basal promoter activities for the three minPs by taking their mean activities when paired with negative controls ORFs and found that the three were highly similar (**Supplementary Fig. 7g**). Comparing activities across all minP sets, we observed a similarly strong positive correlation between promoter and enhancer activities (Spearman’s *ρ* = 0.75, *P*-value < 0.001; **Supplementary Fig. 7h**), suggesting that elements with higher promoter activity also generally tend to exhibit higher enhancer activity irrespective of their paired minP.

In agreement with earlier reports, we observed that minPs differed in their response dynamic range to the same set of elements. For example, the housekeeping minP pGAPDH tended to exhibit a narrower spread of boost indices compared to pMYC or pAPOBEC3F (**Fig. 6a**), consistent with the prior finding that housekeeping promoters displayed the smallest dynamic range^44^. In parallel, our data indicated that differences were not absolute but quantitative, where enhancers that were highly compatible with one promoter generally showed above-baseline activity across all others. To further investigate for potential evidence of enhancer-promoter compatibility, we compared boost indices pairwise across minPs. We found strong correlations in enhancer activity across minP libraries (Spearman’s *ρ* = 0.69 for pMYC vs. pGAPDH, *ρ* = 0.69 for pAPOBEC3F vs. pGAPDH, and *ρ* = 0.78 for pAPOBEC3F vs. pMYC, all *P*-value < 0.001; **Supplementary Fig. 8a**). These results suggest that enhancer-promoter compatibility is broadly conserved, but with minP-specific quantitative shifts that modulate the overall dynamic range. Together, our observations echo the general principle of promoter-dependent responsiveness described previously^43,45^, finding that enhancer function is not strictly promoter-specific but instead quantitatively tuned by promoter context.

**Fig. 6:**
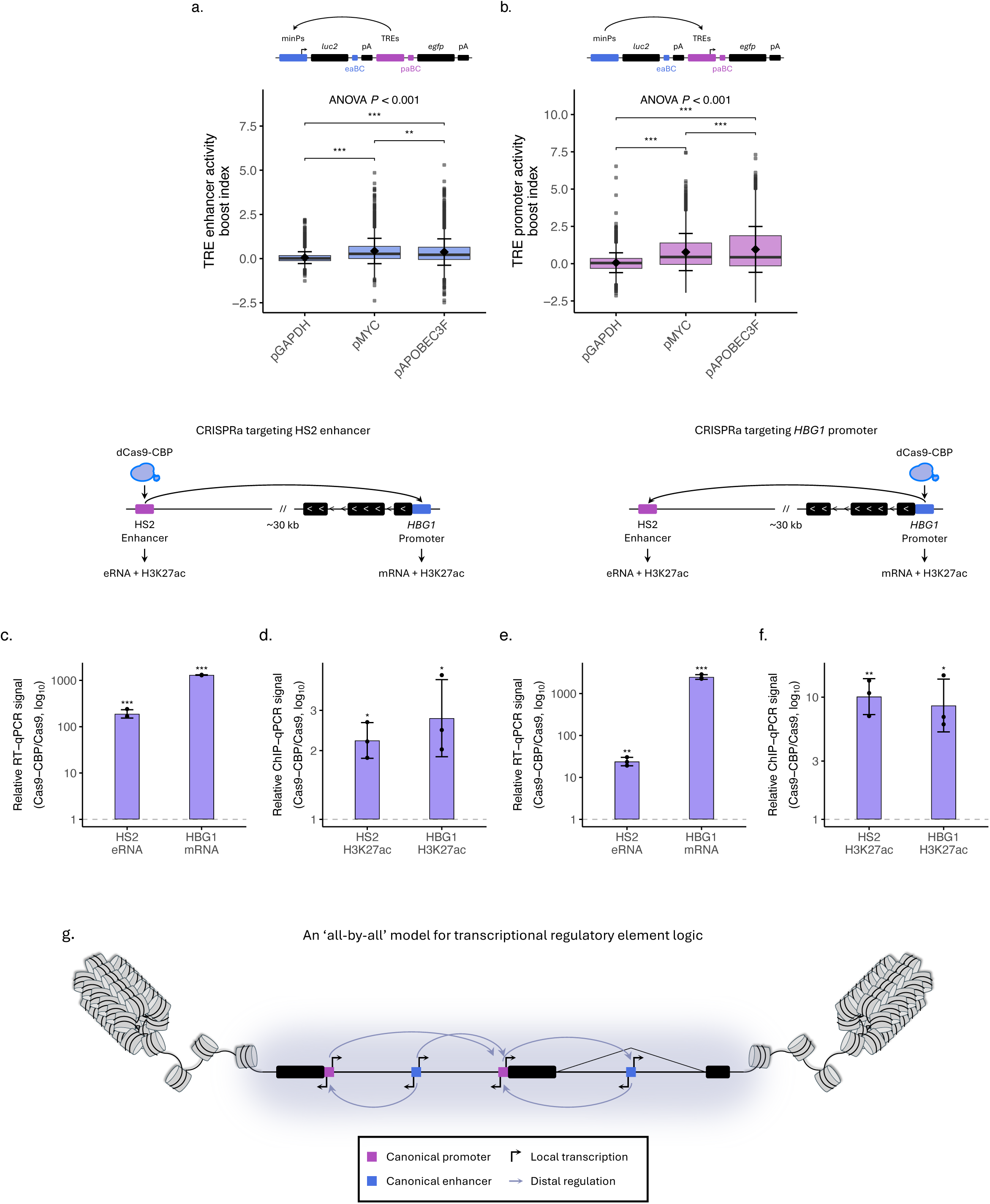
Paired elements exert reciprocal regulatory influences to establish activities of both elements. **a**-**b**, TRE enhancer (a) and promoter (b) activity boost index when paired with the minPs pGAPDH, pMYC, and pAPOBEC3F (Enhancer activity, F(2, 8296) = 309.5, *P*-value < 0.001, ANOVA; Promoter activity, F(2, 8301) = 457.7, *P*-value < 0.001; post-hoc analysis: all except pMYC vs. pAPOBEC3F for enhancer activity, *P*-value < 0.001, Tukey’s HSD; pMYC vs. pAPOBEC3F for enhancer activity, *P*-value < 0.001). **c-d**, RT-qPCR measured HS2 eRNA and HBG1 mRNA signals (c) and ChIP-qPCR measured HS2 H3K27ac and HBG1 H3K27ac signals (d), when HS2 enhancer is targeted by dCas9-CBP. **e-f**, RT-qPCR measured HS2 eRNA and HBG1 mRNA signals (e) and ChIP-qPCR measured HS2 H3K27ac and HBG1 H3K27ac signals (f), when HBG1 promoter is targeted by dCas9-CBP. **g**, An ‘all-by-all’ model for transcriptional regulatory logic. Two promoters (purple) and two enhancers (blue) are shown. Black blocks indicate exons. Diagonal lines indicate splice sites, used to denote intergenic and intragenic enhancers. Black arrows denote promoter activity, lavender arrows enhancer activity. Promoters and enhancers can exhibit functional duality. To exhibit dual functionality, promoter activity is necessary, but not sufficient for enhancer function. Promoters and enhancers can also exert regulatory reciprocity. Certain limitations, such as element compatibility or competition, may restrict associations and thus reciprocal regulatory modulation. Cartoon adapted from Andersson & Sandelin^1^ and Lenhard, Sandelin, & Carninci^3^. For box plots, center line represents median, while box limits indicate upper and lower quartiles. Whiskers extend to 1.5 × interquartile range, and points beyond whiskers denote outliers. Black diamonds represent mean activity, with error bars indicating ±1 standard deviation. Significance was assessed using ANOVA followed by Tukey’s post hoc test. For bar plots, bars show mean RT-qPCR signal (Cas9-CBP relative to Cas9), error bars ±1 SD of log_10_-transformed values. Points represent individual biological replicates. Significance was assessed using One-sample t-test on log_10_ values. Significance levels indicated by asterisks (* *P*-value < 0.05, ** *P*-value < 0.01, *** *P*-value < 0.001). CRISPRa data from Wang et al.^32^

Next, to gain insight into the impact that different minPs have on both the promoter and enhancer activities of the same element, we calculated the promoter and enhancer activity boost indices for each minP library (see **Methods**). Interestingly, pairing with different minPs had highly significant effects on both the promoter and enhancer activities of tested elements (Promoter activity, F(2, 8301) = 457.7, *P*-value < 0.001, ANOVA; Enhancer activity, F(2, 8296) = 309.5, *P*-value < 0.001; **Fig. 6a-b**). Post-hoc analysis revealed that elements paired with pMYC and pAPOBEC3F drove significantly higher promoter activity than when paired with pGAPDH, with a mean difference between pMYC and pGAPDH of 0.71 (*P*-value < 0.001, Tukey’s HSD; **Fig. 6b**), pAPOBEC3F and pGAPDH showing an even greater difference of 0.89 (*P*-value < 0.001), and a mean difference between pMYC and pAPOBEC3F showing a smaller but still significant effect of 0.18 (*P*-value < 0.001). Similarly, Tukey’s HSD revealed that elements paired with pMYC and pAPOBEC3F exerted significantly higher enhancer activity than when paired with pGAPDH, with mean differences of 0.38 (*P*-value < 0.001; **Fig. 6a**) and 0.32 (*P*-value < 0.001), respectively, and with a difference between pMYC and pAPOBEC3F smaller at -0.06 (*P*-value < 0.01). These data suggest that reciprocal regulatory influences between associated elements together establish their respective functions.

### CRISPRa validation of reciprocal activation in the human *β*-globin locus

To independently validate reciprocal regulation in an endogenous genomic context, we re-analyzed a published CRISPRa dataset targeting the human HS2-HBG1 enhancer-promoter pair^32^. The HS2-HBG1 enhancer-promoter pair is a well-established model for studying tissue-specific gene regulation^46^. The *β*-globin locus comprises five *β*-like globin genes (*HBE1*, *HBG1*, *HBG2*, *HBD*, and *HBB*), which are developmentally regulated by a shared upstream locus control region (LCR) enhancer cluster. This LCR contains five discrete DNase I hypersensitive sites (HS1-HS5), among which HS2 functions as a potent erythroid-specific enhancer. Due to its robust enhancer activity and well-defined regulatory interactions, the HS2-HBG1 pair provides a canonical system for dissecting enhancer-promoter communication and transcriptional activation in their native genomic context.

As expected, targeting of the HS2 enhancer with dCas9-CBP increased HS2 eRNA (*P*-value < 0.001, One-sample t-test; **Fig. 6c**) and H3K27ac (*P*-value < 0.05; **Fig. 6d**), in addition to HBG1 mRNA (*P*-value < 0.001; **Fig. 6c**) and H3K27ac (*P*-value < 0.05; **Fig. 6d**), measured via RT-qPCR and ChIP-qPCR, respectively. Strikingly, targeting the *HBG1* promoter also elevated not only HBG1 mRNA (*P*-value < 0.001; **Fig. 6e**) and H3K27ac (*P*-value < 0.05; **Fig. 6f**) but also HS2 eRNA (*P*-value < 0.01; **Fig. 6e**) and H3K27ac (*P*-value < 0.01; **Fig. 6f**). These results provide direct evidence in a native genomic context that promoters can reciprocally enhance the promoter activity of their enhancers, consistent with the regulatory reciprocity observed in our QUASARR-seq data.

Taken together, these data reveal a largely overlooked, though likely pervasive phenomenon in transcriptional regulation (**Fig. 6g**). Not only is the promoter activity of the target element (i.e., promoter) influenced by its associated elements (i.e., enhancers), but the promoter activity of the enhancers are also modulated by the target promoter. This dynamic interplay suggests that promoters and enhancers do not operate as independent unidirectional regulatory modules. Instead, promoters themselves exhibit enhancer-like functions to their enhancers (and likely other promoters), contributing to the system’s overall transcriptional potential, all while enhancers reciprocally influence the functional state of their target promoters (and likely other enhancers).

## Discussion

Our findings further challenge the conventional distinction between promoters and enhancers and instead support a unified model arising from a shared regulatory logic and sequence syntax present across element classes. Using QUASARR-seq, we demonstrate that both canonical promoters and enhancers can drive transcription locally and at a distance under the same regulatory contexts, finding evidence that promoter activity may be necessary, but not sufficient for enhancer function. This intrinsic dual potential is rooted in sequence features and regulatory context.

The strong positive correlation between promoter and enhancer activities exerted by the same element could be indicative of a positive feedback loop, where reciprocal modulation between associated elements reinforces their regulatory functions. This bidirectional feedback may increase and maintain local concentrations of RNAPII, TFs, and co-activators to establish environments with high transcriptional activity. To this end, we propose an ‘all-by-all’ model for transcriptional regulatory elements (**Fig. 6g**), in which all elements (both canonical gene promoters and enhancers) exhibit functional duality to reciprocally regulate one another. This framework extends existing hub/condensate models by adding a bidirectional feedback component, where rather than unidirectional enhancer-to-promoter activation, all elements (i.e., enhancer-to-promoter, promoter-to-enhancer, enhancer-to-enhancer, and promoter-to-promoter) reinforce one another through shared recruitment of factors, sustaining self-organized environments of high transcriptional activity. We note that the concept of regulatory element co-regulation is supported by a recent study^47^. Using single-cell live imaging in flies, Levo et al. elegantly demonstrated that physically distant promoters can undergo coordinated transcriptional bursting. These findings indicate that regulatory crosstalk between TREs may be a widespread feature of gene regulation *in vivo*.

Variants that disrupted promoter activity simultaneously impaired enhancer function, revealing the interconnected nature of these activities. This shared susceptibility has significant implications for understanding disease-associated variants as they may have compounding effects on gene regulation. These findings may underscore the need for therapeutic approaches that address the dual roles of regulatory elements.

While our study provides compelling evidence for a unified framework of TREs, we acknowledge several limitations. Read-through past the first pA accounted for ∼16% of *egfp* signal. Next generation designs could integrate capped RNA enrichment to detect transcripts arising exclusively from the TREs and enable positional TSS mapping, similar to the recently developed TSS-MPRA^48,49^. Moreover, the transient nature of our assay does not fully recapitulate endogenous chromatin dynamics that influence regulatory interactions, in particular the much longer genomic distances spanned by enhancer-promoter contacts. Additionally, broader testing of TREs, minPs, and mutants of the two will be essential to validate and extend our findings and further elucidate the contextual factors that shape functional duality. Future work should move toward *in vivo* models and explore diverse cellular contexts, including shifts with cell identity and induced differentiation. Still, we argue that a reductionist system that can isolate and dissect activities like QUASARR-seq is essential for elucidating fundamental principles of regulatory element logic as complex endogenous systems can be subjected to immense regulatory confounders.

To this end, our study advances our understanding of transcriptional regulation by demonstrating that promoters and enhancers are not separate classes of regulatory elements, as their functional convergence is underpinned by a shared regulatory logic and sequence syntax. This paradigm shift provides new insights into the mechanisms of gene regulation and the functional consequences of regulatory variants in health and disease.

## Materials and Methods

### TRE definition and selection

To classify promoter and enhancer elements, we based our approach solely on transcriptional activity (PRO-cap), chromatin accessibility (DNase-seq), and proximity to known gene annotations (GENCODE) in K-562, given that not all accessible regions are transcriptionally active and that histone signatures traditionally used to distinguish promoters and enhancers converge when local transcription is considered.

Thus, to systematically compare promoter and enhancer elements, we defined candidate active promoters as proximal PRO-cap TSSs (distance cutoff < 500 bp from GENCODE annotation), and candidate active enhancers as distal PRO-cap TSSs (distance cutoff > 500 bp from GENCODE annotation). Similarly, we defined candidate inactive promoters as proximal PRO-cap untranscribed DNase-seq DHSs (distance cutoff < 500 bp from GENCODE annotation), and candidate inactive enhancers as distal PRO-cap untranscribed DNase-seq DHSs (distance cutoff > 500 bp from GENCODE annotation). Transcribed elements were cloned using boundaries set at 60 bp downstream from each divergent TSS peak maximum. Untranscribed elements were cloned using boundaries set at DHS peak coordinates.

As negative controls, we included sequence-verified, TSS/DHS-screened human ORFs that showed no regulatory activity in eSTARR-seq^27^. As positive controls, we included a set of viral promoters/enhancers (*Cytomegalovirus* (CMV) and Rous sarcoma virus (RSV)), HS005^50^, and MYC E1-7^51^.

### TRE cloning

The cloning of TREs was carried out as previously described^27^. In brief, primers were designed in batch using our in-house web tool, which flanks forward primers with attB1′ and reverse primers with attB2′ 5′ overhangs. Primers used for TRE cloning were synthesized by Eurofins. K-562 genomic DNA (E493; Promega Corp.) was used as template for PCR amplifications using Phusion High-Fidelity DNA Polymerase (M0530; New England Biolabs) and PrimeSTAR GXL DNA Polymerase (R050A; Takara Bio Inc.). Amplicons were inserted into pDONR223 via Gateway BP cloning and single-colony-derived entry clones were sequence verified as previously described. Sequence-verified clones were propagated in lysogeny broth (LB) supplemented with spectinomycin, pooled, and purified using E.Z.N.A. Plasmid DNA Midi Kit (D6904; Omega Bio-tek, Inc.).

TREs were inserted into pDEST-luc-pMYC-preBC and pDEST-luc-pMYC-preBC-CCW via *en masse* Gateway LR cloning, propagated in LB supplemented with ampicillin and extracted using E.Z.N.A. Endo-Free Plasmid DNA Maxi Kit (D6926; Omega Bio-tek, Inc.). The resulting libraries were used as templates for downstream TRE barcoding (see eaBC-TRE-paBC index generation section).

### QUASARR-seq vector design and engineering

The QUASARR-seq vector design features an interchangeable minP that drives the expression of a reporter gene (*luc2*) that contains a degenerate 20 bp enhancer activity barcode (eaBC) in its 3′ UTR. Candidate TREs are cloned downstream of this reporter system such that they can drive the expression of a second reporter gene (*egfp*) that contains another degenerate 20 bp promoter activity barcode (paBC) in its 5′ UTR. In this design, TREs are positioned 2,180 bp downstream of the minPs (or 2,916 bp upstream when traversing the circular backbone), ensuring a fixed distance across all constructs. Importantly, each reporter system is insulated with a cleavage and polyadenylation site (pA) to avoid confounding activity readouts from spurious initiation from the other element (i.e., initiation at the minP does not confound initiation at the TRE and initiation at the TRE does not confound initiation at the minP).

The QUASARR-seq assay vector, pDEST-QUASARR-luc-gfp-pMYC, was generated by modifying pDEST-luc-pMYC-preBC. To engineer pDEST-QUASARR-luc-gfp-pMYC, an egfp-pA reporter sequence was cloned downstream of the attR1-attR2 Gateway cloning cassette. We also generated the assay vectors pDEST-QUASARR-luc-gfp-pGAPDH, pDEST-QUASARR-luc-gfp-pAPOBEC3F, and pDEST-QUASARR-luc-gfp-pADGRG5 that are identical to pDEST-QUASARR-luc-gfp-pMYC except that the *MYC* minP was replaced with a *GAPDH*, *APOBEC3F*, or *ADGRG5* minP, respectively. minPs were cloned using boundaries set at 60 bp downstream from each PRO-cap-detected divergent TSS peak maximum.

pDEST-luc-pMYC-preBC was generated by modifying our eSTARR-seq assay vector, pDEST-hSTARR-luc-pMYC. To engineer pDEST-luc-pMYC-preBC, the attR1-attR2 cassette in pDEST-hSTARR-luc-pMYC was removed from the 3′ UTR and re-cloned downstream of the pA. To engineer pDEST-luc-pMYC-preBC-CCW, the attR1-attR2 cassette in pDEST-luc-pMYC-preBC was removed and then re-cloned back into its original position in reverse orientation.

The vectors pDEST-luc-pMYC-preBC and pDEST-luc-pMYC-preBC-CCW were used as surrogates to clone TREs in forward and reverse orientation, respectively and were used as templates for generating eaBC-TRE-paBC amplicons, which were subsequently cloned into the QUASARR-seq assay vector.

### eaBC-TRE-paBC index generation

Generation of high-complexity eaBC-TRE-paBC indices are required for accurate and robust enhancer and promoter activity measurements. High complexity maps were achieved through the generation of eaBC-TRE-paBC amplicons via consecutive isothermal oligo extensions using Phusion High-Fidelity DNA Polymerase (M0530; New England Biolabs). TREs were first cloned in both forward and reverse orientation into the surrogate vectors pDEST-luc-pMYC-preBC and pDEST-luc-pMYC-preBC-CCW, respectively. These libraries were used as templates for primer extension reactions using oligos that contain 1. 3′ ends complementary to TRE flanking sequences of the surrogate vectors, 2. internal 20 bp degenerate sequences, and 3. 5′ ends homologous to the QUASARR-seq assay vectors. Amplicons were cloned into pDEST-QUASARR-luc-gfp using NEBuilder HiFi DNA Assembly Master Mix (E2621; New England Biolabs), transformed, and plated on LB/agar supplemented with ampicillin. Colonies were scrapped into LB and extracted using E.Z.N.A. Endo-Free Plasmid DNA Maxi Kit (D6926; Omega Bio-tek, Inc.).

To ensure activity score accuracy and robustness and reduce potential eaBC/paBC-specific biases, a minimum of 10 eaBCs/paBCs should be uniquely assigned to each TRE. Thus, to obtain adequate eaBC-TRE-paBC coverage, the number of colonies required should exceed the product of the number of TREs being surveyed and the desired number of unique eaBCs/paBCs (i.e., surveying 100 TREs each with 100 eaBCs/paBCs requires a minimum of 10,000 colonies). To evaluate input library complexity and construct an eaBC-TRE-paBC index, pre-transfected eaBC-TRE-paBC libraries were generated prior to electroporation.

### Cell culture and nucleofection

K-562 (CCL-243; ATCC) cells were cultured in IMDM (30-2005; ATCC) supplemented with 10% FBS (30-2020; ATCC) at 37°C with 5% CO_2_. QUASARR-seq input libraries were electroporated into K-562 using Amaxa Cell Line Nucleofector Kit V (VCA-1003; Lonza Group AG) with a Lonza Amaxa Nucleofector II Device using program T-016. The manufacturers’ Amaxa Cell Line Nucleofector Kit V protocol for ATCC K-562 was followed, except for each electroporation, 1 x 10^6^ cells received 20 μg of QUASARR-seq library. Five electroporations were carried out per technical replicate, three technical replicates per biological replicate. Cells for different biological replicates were cultured and electroporated on separate days.

Following a 6-hour incubation period, cells were harvested by pooling samples for the same technical replicate to remove variance introduced during electroporation. Following three washes with 1X PBS (10010023; Gibco), each technical replicate was parsed out where 4 x 10^6^ cells were used for RNA libraries and 1 x 10^6^ cells were used for DNA libraries. Cell pellets were snap-frozen with LN_2_ and stored in -80°C until library preparation.

### Optimal harvest time experiments

To determine the optimal post-transfection harvest time for plasmid-derived RNA expression, we performed a series of time-course experiments in K-562 as part of our previous study^27^. As a pilot experiment, 1 x 10^6^ cells were electroporated with 10 μg of pDEST-hSTARR-luc-pMYC-RSV (10 pg/cell) using the Amaxa Cell Line Nucleofector Kit V with a Lonza Amaxa Nucleofector II Device using program T-016. At 6, 12, 24, 36, and 48 hours post transfection (hpt), 5 x 10^5^ cells were harvested per time point. Total RNA was extracted with TRIzol Reagent (15596026; Invitrogen), treated with DNase I (AM2222; Invitrogen), and purified using RNeasy Mini Kit (74104; QIAGEN). Reverse transcription was performed using an Oligo d(T)_18_ mRNA Primer (S1316S; New England Biolabs), and cDNA was quantified by qPCR using exon junction-spanning primers specific for plasmid-derived RNA and for GAPDH mRNA as an internal control. Technical triplicates (n = 3) were averaged per time point. As a follow up experiment, cells were electroporated with pDEST-hSTARR-luc-pMYC-Control1K or pDEST-hSTARR-luc-TRE202, and 4 x 10^5^ cells were collected at 2, 4, 6, 12, 24, 36, and 48 hpt. RNA isolation, DNase treatment, purification, reverse transcription, and qPCR were performed as in the pilot experiment. As a final experiment, cells were electroporated with pDEST-hSTARR-luc-TRE202 or pDEST-hSTARR-luc-pMYC-TRE202, and 4 x 10^5^ cells were harvested at 2, 4, 6, 12, and 24 hpt. Processing and analysis were performed as in the prior experiments.

Across all experiments, plasmid-derived transcript abundance consistently peaked at 6 hpt and declined or plateaued at later time points. Based on these observations, 6 hpt was selected as the standard harvest point for subsequent assays (**Supplementary Fig. 9a-c**).

### QUASARR-seq library preparation

QUASARR-seq requires a minimum of four libraries per technical replicate: two RNA libraries targeting the eaBC and paBC transcripts, respectively, and two DNA libraries targeting the eaBC and paBC inputs, respectively. Total RNA was extracted from cells using TRIzol Reagent following the manufacturer’s instructions. To remove contaminating plasmid DNA, RNA was treated with DNase I and subsequently purified using RNeasy Mini Kit. Reverse transcription was performed with the DNase-treated RNA as the template using SuperScript III Reverse Transcriptase (18080093; Invitrogen). Plasmid DNA was extracted from cells as previously described^52^. A first primer extension was performed with the extracted DNA as the template. Reactions were treated with Exonuclease I (M0293; New England Biolabs) to remove unused primer and purified using DNA Clean & Concentrator-5 (D4013; Zymo Research Corp.). A second primer extension was performed with the products of the reverse transcription (RNA libraries) and the first primer extension (DNA libraries) as the templates, respectively. Reactions were again treated with Exonuclease I and purified using DNA Clean & Concentrator-25 (D4033; Zymo Research) and DNA Clean & Concentrator-5 for the RNA and DNA libraires, respectively. Finally, low-cycle PCR was performed to add sequencing adapters, followed by acquisition of 2 × 150 bp reads on an Illumina NovaSeq X Plus. All primer sequences used in this work can be found in **Supplementary Table 1**.

### RT-qPCR

Cell culture and nucleofection were carried out as described above, except 1 x 10^6^ cells were electroporated with 5 µg plasmid DNA and incubated for 24 h. Total RNA was extracted as described above. cDNA was generated from 1 µg of DNase-treated RNA using LunaScript RT SuperMix Kit (E3010; New England Biolabs) and Random Primer Mix (S1330S; New England Biolabs). Parallel no-RT controls were performed to assess plasmid DNA contamination. cDNA was purified with DNA Clean & Concentrator-25. qPCR was carried out using Luna Universal qPCR Master Mix (M3003; New England Biolabs) with gene-specific primers targeting *luc2* and *egfp* on a QuantStudio 7 Pro Real-Time PCR System (A43183; Applied Biosystems). Amplification was performed with an initial denaturation at 95 °C for 1 min, followed by 40 cycles of 95 °C for 15 s and 60 °C for 30 s.

### RT-qPCR data processing and read-through transcription estimation

For each element and biological replicate, *C*_*q*_ values from technical replicates were averaged. Reporter values (*luc2* and *egfp*) were then aligned per replicate, and relative expression was calculated using a Δ*C*_*q*_ method, where Δ*C*_*q*_ was defined as:

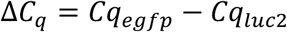

Relative quantification (*RQ*) was computed as:

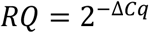

For each element, mean *RQ*, standard deviation (*SD*), and the standard error of the mean (*SEM*) were calculated across replicates.

To estimate read-through transcription, mean *RQ* values for the empty vector control (*RQ*_*Empty*_) and TRE-driven positives (*RQ*_*TRE*_) were compared. The fraction of *egfp* signal attributable to read-through was calculated as:

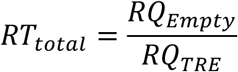

### QUASARR-seq data preprocessing

QUASARR-seq data preprocessing includes two main parts: 1. eaBC-TRE-paBC index mapping and 2. RNA aBC/DNA aBC activity measurement. Raw sequencing data was first filtered using fastp using the following parameters "--disable_adapter_trimming --trim_poly_g --cut_right", followed by processing using biodatatools, with detailed commands and workflow found in the **Extended Text**. Briefly, sequence information was retrieved according to the library’s corresponding layout; eaBC-TRE-paBC and unique molecular identifiers (UMIs) for index mapping, and RNA aBCs, DNA aBCs, and UMIs for activity measurements.

In index mapping, eaBCs and paBCs were clustered by merging all sequences with a maximum Hamming distance of one for sequencing error tolerance. Partial element sequences were aligned to the reference element sequences. eaBCs and paBCs were then assigned to the TREs. For the multiple minP library set, an additional three-bp minP barcode was used to assign aBCs to their associated minPs. eaBCs and paBCs were regarded as a representation of a given TRE(A), if the number of eaBC-TRE(A)-paBC entries were significantly higher than all other eaBC-TRE(X)-paBC entries for any given TRE(X). In activity measurement, eaBCs and paBCs were extracted and matched to the previously clustered eaBCs and paBCs during index mapping and assigned to their corresponding TRE. All counts generated were subjected to UMI correction to collapse identical copies. The resultant RNA and DNA element counts were then normalized using edgeR to calculate the normalized element counts and logFCs.

### Activity boost index calculation

Activity boost index:

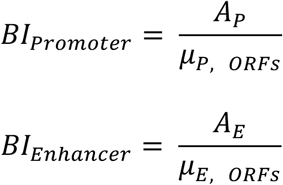

Where:

*A*_*P*_ and *A*_*E*_ are the measured promoter and enhancer activity logFCs, respectively.

*μ*_*P, ORFS*_ and *μ*_*E, ORFS*_ are the mean promoter and enhancer activities of negative control ORFs, respectively.

### Uniform active call pipeline

To identify active TREs, we applied a uniform active element call pipeline, as previously described^27^. The pipeline begins with raw count matrices for DNA and RNA libraries. A pre-filtering procedure retains TREs that have counts per million (CPM) greater than a threshold calculated based on a raw count of 10 in the smallest DNA library. To account for library size differences and composition biases, a modified version of the trimmed mean of M values (TMM) normalization method^53^ was utilized to rely solely on negative control ORFs to normalize across libraries. log_2_-transformed RNA-to-DNA ratios (log_2_(RNA aBCs/DNA aBCs)) were computed as a measure of regulatory activity for each TRE using the limma-voom pipeline^54^. To assess enhancer activity, a Z-score approach was applied to compare the log_2_(RNA eaBCs /DNA eaBCs) values of each TRE to those of negative controls. TREs that have significantly higher regulatory activity than the basal transcription level defined by the negative controls in both orientations were identified as active enhancers. A similar approach was applied to identify active promoters.

### Activity balance index calculation

Z-score normalization:

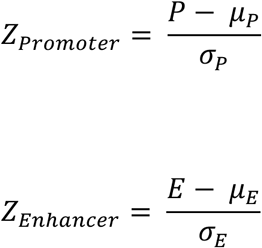

Where:

*P* and *E* are the measured promoter and enhancer activity boost indices, respectively.

*μ*_*P*_ and *μ*_*E*_ are the mean activity boost indices for promoter and enhancer, respectively.

*σ*_*P*_ and *σ*_*E*_ are the standard deviations of the promoter and enhancer activity boost indices, respectively.

Activity balance index calculation:

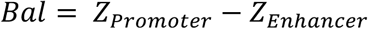

Where *Bal* is the activity balance index. Functional classification:

Promoter-dominant: *Bal* > 1

Enhancer-dominant: *Bal* < −1

Balanced: −1 ≤ *Bal* ≤ 1

### GC content calculation

GC content calculation:

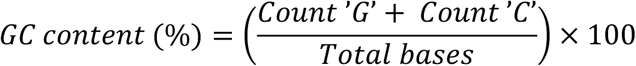

Where:

*Count* ’*G*’ and ’*C*’ is the total number of guanine and cytosine bases in the sequence and *Total bases* is the total length of the sequence (including all bases A, T, G, and C).

### Motif match score calculation

For a given sequence *x*, the motif match score, *match*(*x*), was calculated as the log-odds ratio between the probability of observing the sequence under a position weight matrix (PWM) distribution, and the probability of observing the sequence under a background model:

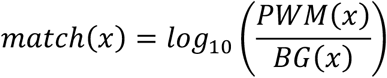

Where:

The probability under the PWM distribution, *PWM*(*x*), was determined by the product of the probabilities of each base in the sequence based on the PWM:

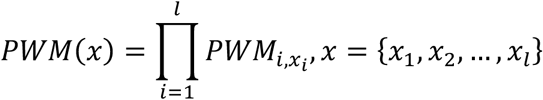

PWMs used in this calculation were obtained from Adato & Sloutskin et al.^55^

The probability under the background distribution, *BG*(*x*), was calculated as the product of the fractions of each base observed across all GRO-cap peaks in K-562 using data from Core & Martins et al.^14^

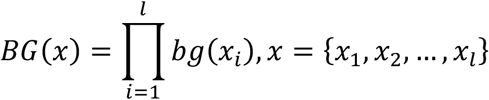

Where:

The background probabilities, *bg*(*x*_*i*_) for each base were defined as:

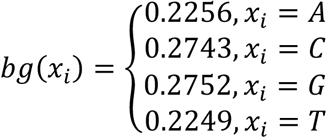

### Change boost index calculation

Δboost index:

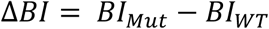

Where:

*BI*_*WT*_ and *BI*_*Mut*_ are the wild-type and mutant boost indices, respectively.

### PRO-cap/seq data analysis

PRO-cap signal was defined as the total number of UMI-deduplicated read counts within the boundaries of tested elements. PRO-seq signal was defined as either 1, the total number of UMI-deduplicated read counts 100 (**Supplementary Fig. 2a**), 250 (**Supplementary Fig. 2b**), or 500 (**Supplementary Fig. 2c**) bp downstream (for forward elements) and upstream (for reverse elements) from the boundaries of tested elements, 2, the mean number of UMI-deduplicated read counts 100 (**Supplementary Fig. 2d**), 250 (**Supplementary Fig. 2e**), or 500 (**Supplementary Fig. 2f**) bp downstream and upstream from the boundaries of tested elements, 3, the total number of UMI-deduplicated read counts within the boundaries of tested elements including a 100 (**Supplementary Fig. 2g**), 250 (**Supplementary Fig. 2h**), or 500 (**Supplementary Fig. 2i**) bp extension downstream (for forward elements) and upstream (for reverse elements) from the boundaries of tested elements, or 4, the mean number of UMI-deduplicated read counts within the boundaries of tested elements including a 100 (**Supplementary Fig. 2j**), 250 (**Supplementary Fig. 2k**), or 500 (**Supplementary Fig. 2l**) bp extension downstream and upstream from the boundaries of tested elements.

### SuRE, lentiMPRA, ATAC-STARR-seq, and WHG-STARR-seq data analysis

To benchmark QUASARR-seq against orthogonal assays, we compared elements with at least a 10% reciprocal overlap also tested in SuRE, and at least a 50% reciprocal overlap also tested in lentiMPRA, ATAC-STARR-seq, and WHG-STARR-seq. Active enhancer calls for lentiMPRA, ATAC-STARR-seq, and WHG-STARR-seq were obtained using the same uniform active element call pipeline as described.

## Data availability

QUASARR-seq data will be made available in the ENCODE portal (www.encodeproject.org). PRO-cap (accession no. ENCSR220XSM) data were retrieved from the ENCODE portal. PRO-seq data will be made available in the ENCODE portal. Processed SuRE data were obtained from van Arensbergen et al.^29^ lentiMPRA (accession no. ENCSR382BVV), ATAC-STARR-seq (accession no. ENCSR312UQM), and WHG-STARR-seq (accession no. ENCSR661FOW) data were retrieved from the ENCODE portal. Processed CRISPRa RT- and ChIP-qPCR data from Wang et al.^32^ were obtained from I. Hilton, Rice University. All accession codes not provided will be made available prior to publication.

## Code availability

Code will be made available on GitHub prior to publication.

## Acknowledgements

We thank I. Hilton for sharing the CRISPRa datasets from Wang et al.^32^ This work was supported by the National Institutes of Health grants R01 AG077899 and R01 HG012970 to J.T.L. and H.Y. M.I.P was supported in part by a Cornell University Center for Vertebrate Genomics Scholarship.

## Author contributions

M.I.P. conceptualized the study with guidance from S.R.S., H.Y., and J.T.L. M.I.P. conceptualized, designed, and developed QUASARR-seq with input from S.R.S. M.I.P. generated the QUASARR-seq and SOLARR-seq data. A.K-Y.L. developed the QUASARR-seq data pre-processing pipeline. A.K-Y.L. performed the QUASARR-seq data pre-processing. J.Z. developed the uniform activity calling pipeline. M.I.P. performed the downstream data analysis. M.I.P. generated the figures with guidance from H.Y. M.I.P. wrote the manuscript with guidance from H.Y., J.T.L., A.O., and S.R.S. N.D.T. designed the TRE selection criterion. J.L. cloned the TREs. L.Y. designed the mutant TREs. Y.J. cloned the minPs. X.P. performed PRO-seq data curation and motif match score calculations. H.Y. and J.T.L. oversaw the project. H.Y. and J.T.L. acquired funding.

