## Supplementary Figures 1-9 for "Simultaneous measurement of intrinsic promoter and enhancer potential reveals principles of functional duality and regulatory reciprocity"

a.

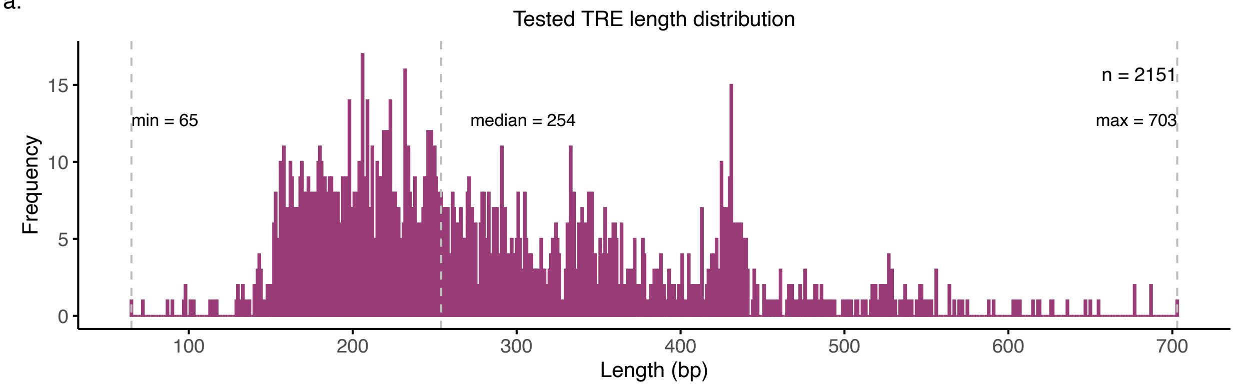

Tested TRE length distribution
n = 2151
min = 65
median = 254
max = 703
Frequency
0
5
10
15
100
200
300
400
500
600
700
Length (bp)

**Supplementary Fig. 1: Length distribution of tested TREs.**
**a**, A total of 2,151 TREs were tested, ranging from 65 to 703 bp, with a median length of 254 bp.

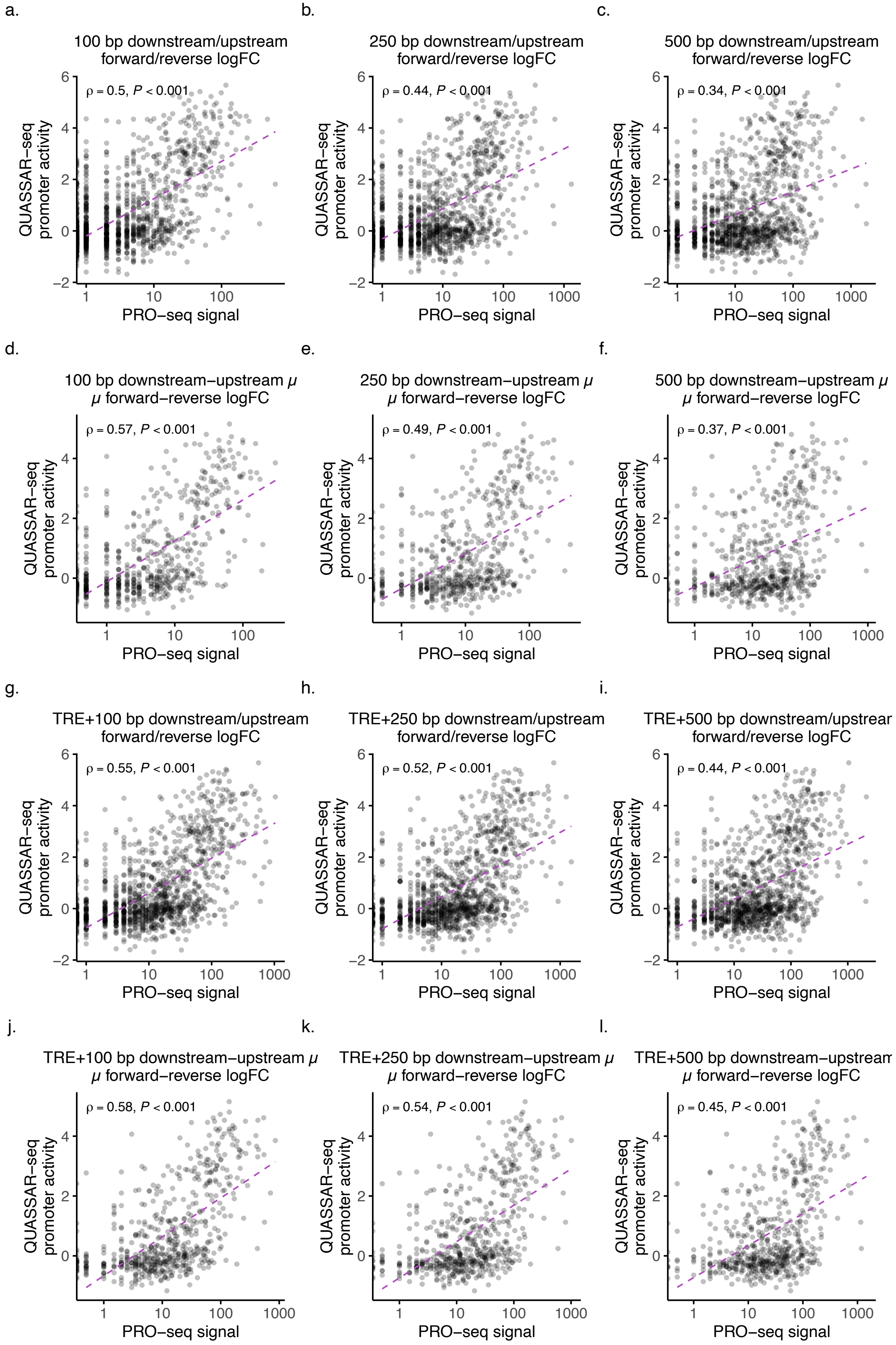

a.
100 bp downstream/upstream forward/reverse logFC
ρ = 0.5, P < 0.001
b.
250 bp downstream/upstream forward/reverse logFC
ρ = 0.44, P < 0.001
c.
500 bp downstream/upstream forward/reverse logFC
ρ = 0.34, P < 0.001
d.
100 bp downstream−upstream μ μ forward−reverse logFC
ρ = 0.57, P < 0.001
e.
250 bp downstream−upstream μ μ forward−reverse logFC
ρ = 0.49, P < 0.001
f.
500 bp downstream−upstream μ μ forward−reverse logFC
ρ = 0.37, P < 0.001
g.
TRE+100 bp downstream/upstream forward/reverse logFC
ρ = 0.55, P < 0.001
h.
TRE+250 bp downstream/upstream forward/reverse logFC
ρ = 0.52, P < 0.001
i.
TRE+500 bp downstream/upstrear forward/reverse logFC
ρ = 0.44, P < 0.001
j.
TRE+100 bp downstream−upstream μ μ forward−reverse logFC
ρ = 0.58, P < 0.001
k.
TRE+250 bp downstream−upstream μ μ forward−reverse logFC
ρ = 0.54, P < 0.001
l.
TRE+500 bp downstream−upstrean μ forward−reverse logFC
ρ = 0.45, P < 0.001
QUASSAR−seq promoter activity
PRO−seq signal

**Supplementary Fig. 2: QUASARR-seq promoter activity comparisons with PRO-seq signal. a-c**, Correlation between QUASARR-seq promoter activity, with elements in forward and reverse orientation treated as separate, and downstream (for forward) and upstream (for reverse) PRO-seq signal 100 (a), 250 (b), or 500 (c) bp from element. **d-f**, Correlation between QUASARR-seq promoter activity, using the mean of elements in forward and reverse orientation, and the mean of downstream and upstream PRO-seq signal 100 (d), 250 (e), or 500 (f) bp from element. **g-i**, Correlation between QUASARR-seq promoter activity, with elements in forward and reverse orientation treated as separate, and PRO-seq signal within the element and a 100 (g), 250 (h), or 500 (i) bp extension downstream (for forward) and upstream (for reverse) from element. **j-l**, Correlation between QUASARR-seq promoter activity, using the mean of elements in forward and reverse orientation, and PRO-seq signal within the element and the mean of downstream and upstream of 100 (j), 250 (k), or 500 (l) bp extensions from element. Spearman's rank correlation coefficient.

a. b.

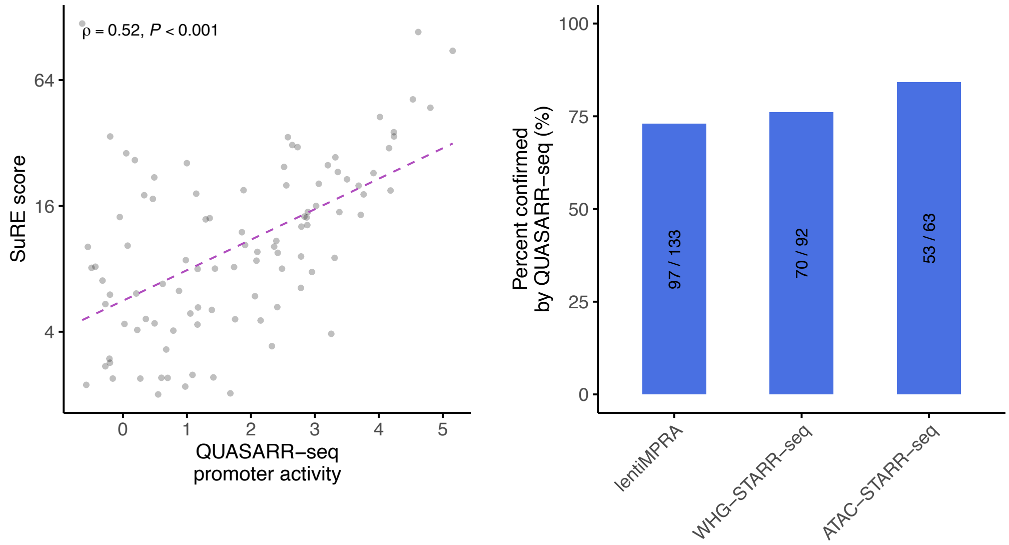

ρ = 0.52, P < 0.001
SuRE score
64
16
4
0
1
2
3
4
5
QUASARR−seq
promoter activity
Percent confirmed
by QUASARR−seq (%)
100
75
50
25
0
97 / 133
70 / 92
53 / 63
lentiMPRA
WHG−STARR−seq
ATAC−STARR−seq

**Supplementary Fig. 3: QUASARR-seq activity benchmarking with orthogonal assays.**
**a**, Correlation between QUASARR-seq promoter activity and Survey of Regulatory Elements (SuRE) score (Spearman's $\rho = 0.52$, $P$-value < 0.001). Calculated based on 10% reciprocal overlap between elements across assays. b, Percent confirmed of lentiMPRA, WHG-STARR-seq, and ATAC-STARR-seq active enhancers by QUASARR-seq. Calculated based on 50% reciprocal overlap between elements across assays.

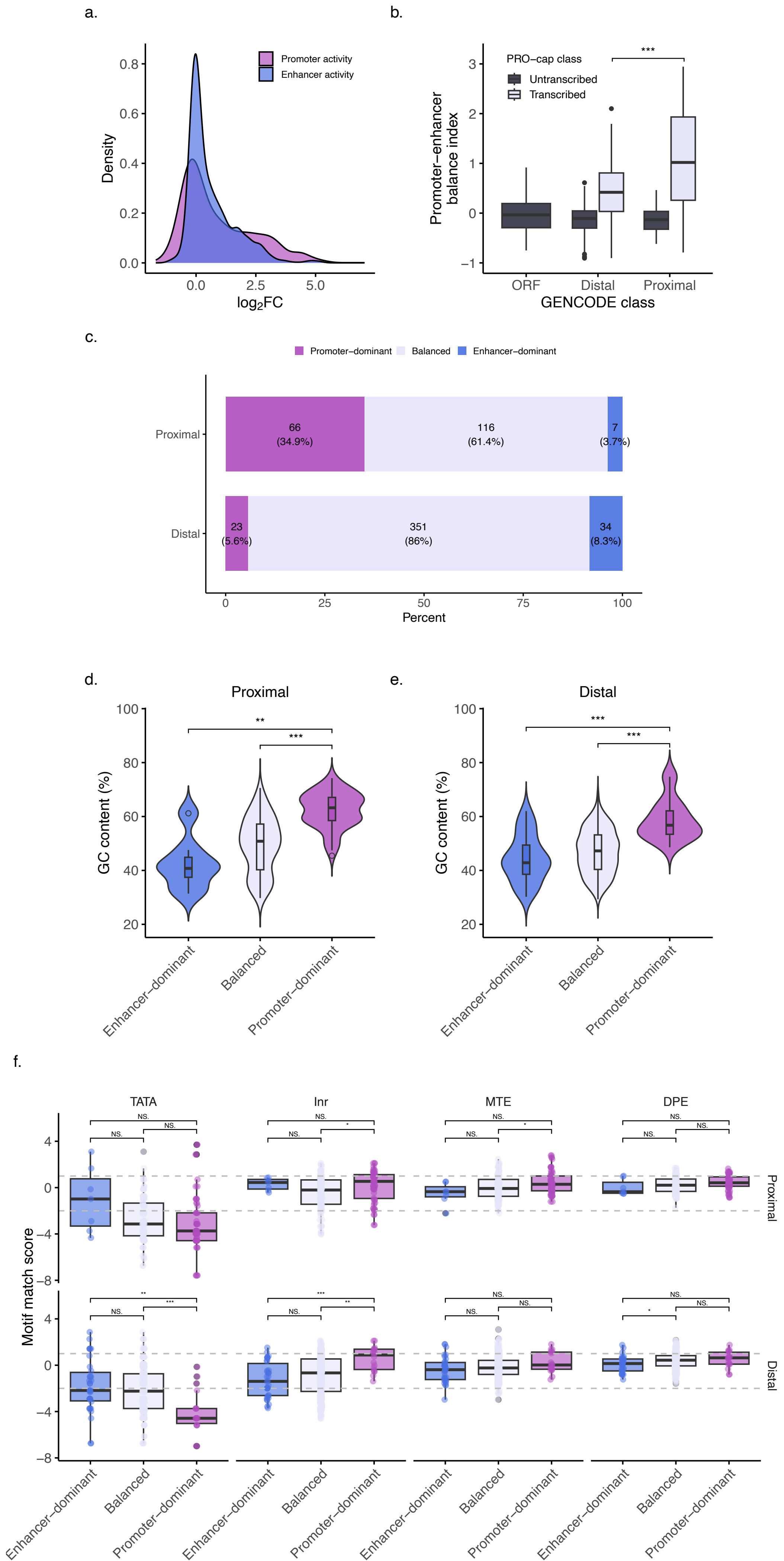
a.
Promoter activity
Enhancer activity
Density
log2FC
b.
PRO−cap class
Untranscribed
Transcribed
Promoter−enhancer balance index
ORF
Distal
Proximal
GENCODE class
***
c.
Promoter−dominant
Balanced
Enhancer−dominant
Proximal
Distal
66 (34.9%)
116 (61.4%)
7 (3.7%)
23 (5.6%)
351 (86%)
34 (8.3%)
Percent
d.
Proximal
e.
Distal
GC content (%)
Enhancer−dominant
Balanced
Promoter−dominant
f.
TATA
Inr
MTE
DPE
Motif match score
Proximal
Distal
NS.

**Supplementary Fig. 4: GC content and Inr drive functional bias in activity balance across GENCODE classes.**

**a**, QUASARR-seq promoter and enhancer $Log_2FC$ distribution. **b**, Ratio of promoter-to-enhancer activity boost index across GENCODE and PRO-cap classes (*P*-value < 0.001, Student's t-test). **c**, Activity balance category distribution by GENOCDE class. **d-e**, GC content (%) of proximal (d) and distal (e) elements parsed by activity balance category (promoter-dominant vs. balanced for proximal and distal, *P*-value < 0.001; promoter-dominant vs. enhancer-dominant for proximal, *P*-value < 0.01; for distal, *P*-value < 0.001, Student's t-test with Bonferroni correction). **f**, Plus-strand motif match scores by activity category and GENCODE class. For proximal, Inr and MTE promoter-dominant vs. balanced (*P*-value < 0.05, Wilcoxon rank-sum test with Bonferroni correction). For distal, Inr promoter-dominant vs. balanced (*P*-value < 0.01) and vs. enhancer-dominant (*P*-value < 0.001), TATA promoter-dominant vs. balanced (*P*-value < 0.001) and vs. enhancer-dominant (*P*-value < 0.01) elements, DPE balanced vs. enhancer-dominant (*P*-value < 0.05). For box plot, center line represents median, while box limits indicate upper and lower quartiles. Whiskers extend to 1.5 × interquartile range, and points beyond whiskers denote outliers. Significance levels indicated by asterisks (* *P*-value < 0.05, ** *P*-value < 0.01, *** *P*-value < 0.001).

a.

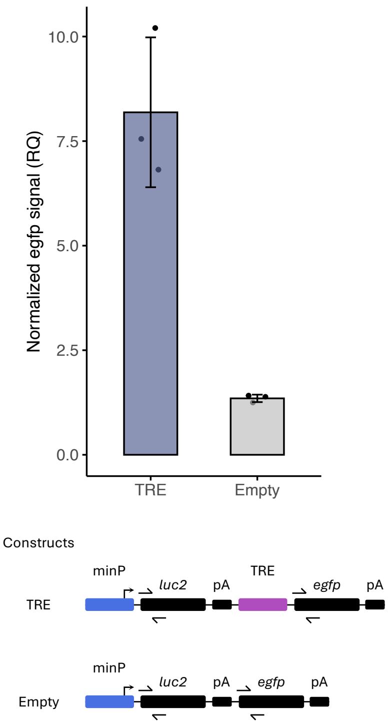
Normalized egfp signal (RQ)
10.0
7.5
5.0
2.5
0.0
TRE
Empty
Constructs
minP
luc2
pA
TRE
egfp
pA
TRE
minP
luc2
pA
egfp
pA
Empty

**Supplementary Fig. 5: Read-through estimation RT-qPCR.**

**a**, Bottom, RT-qPCR targeting *luc2* and *egfp* in TRE-containing and TRE-lacking constructs. Top, *luc2*-normalized *egfp* signal for TRE and empty vectors. Bars show mean RT-qPCR signal, error bars ±1 SD. Points represent individual biological replicates.

a.

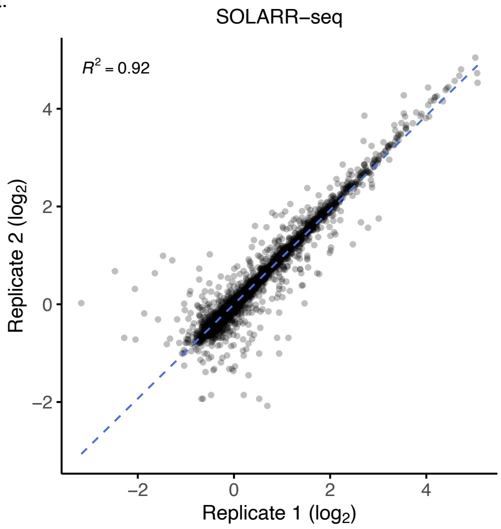
SOLARR−seq
$R^2 = 0.92$
Replicate 2 ($log_2$)
Replicate 1 ($log_2$)
4
2
0
−2
−2
0
2
4

b.

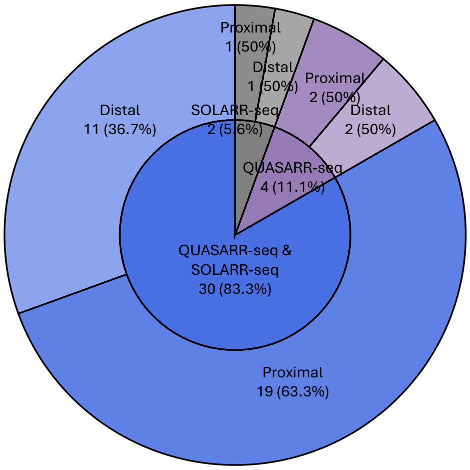
Proximal
1 (50%)
Distal
1 (50%)
Proximal
2 (50%)
Distal
2 (50%)
Distal
11 (36.7%)
SOLARR-seq
2 (5.6%)
QUASARR-seq
4 (11.1%)
QUASARR-seq &
SOLARR-seq
30 (83.3%)
Proximal
19 (63.3%)

**Supplementary Fig. 6: Downstream sequence features do not confer element activity or type.** **a**, Correlation of SOLARR-seq enhancer activity measurements between replicates. **b**, Active enhancer calls by QUASARR-seq and SOLARR-seq parsed by GENCODE class.

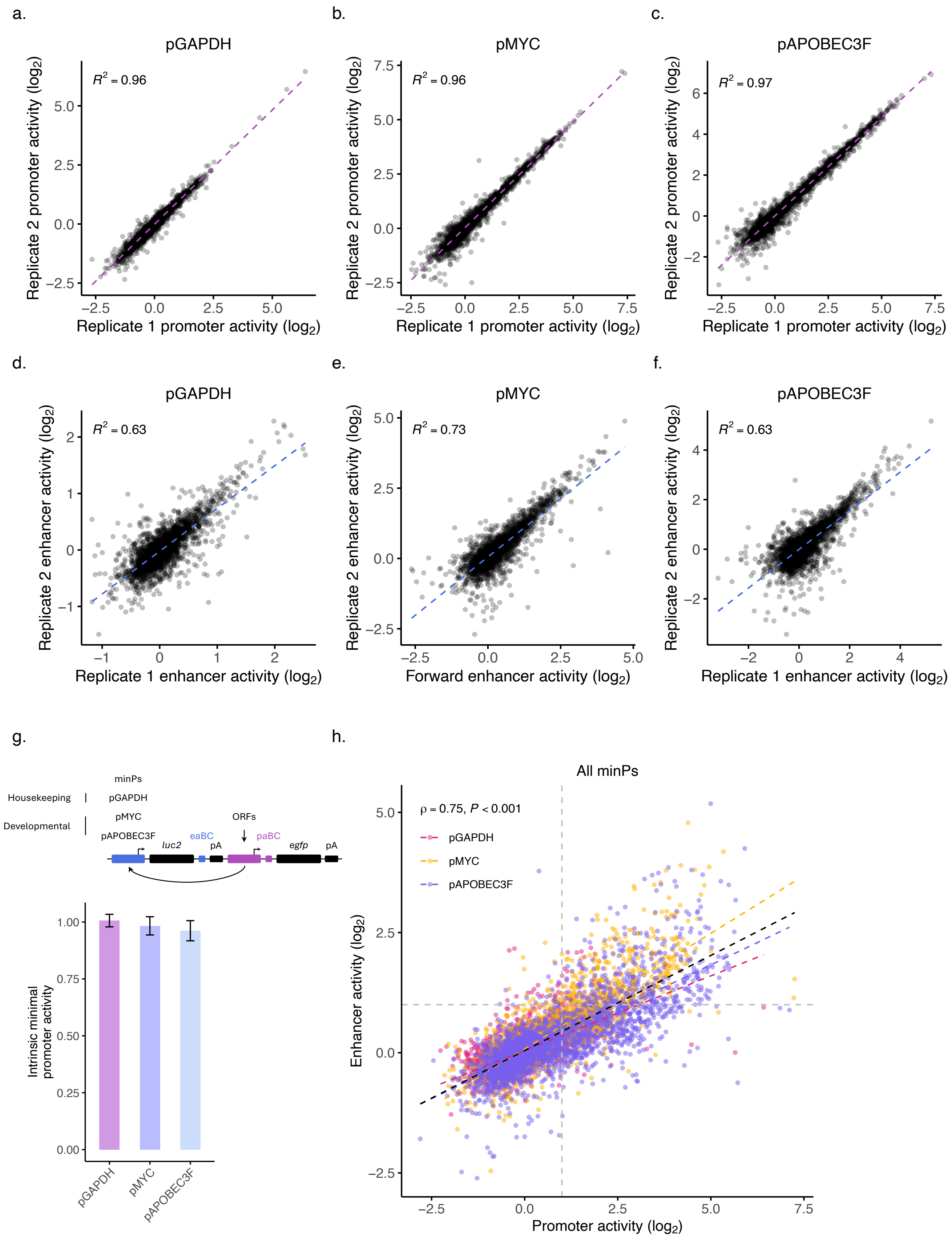

a.
pGAPDH
$R^2 = 0.96$
Replicate 2 promoter activity (log2)
Replicate 1 promoter activity (log2)
b.
pMYC
$R^2 = 0.96$
Replicate 2 promoter activity (log2)
Replicate 1 promoter activity (log2)
c.
pAPOBEC3F
$R^2 = 0.97$
Replicate 2 promoter activity (log2)
Replicate 1 promoter activity (log2)
d.
pGAPDH
$R^2 = 0.63$
Replicate 2 enhancer activity (log2)
Replicate 1 enhancer activity (log2)
e.
pMYC
$R^2 = 0.73$
Replicate 2 enhancer activity (log2)
Forward enhancer activity (log2)
f.
pAPOBEC3F
$R^2 = 0.63$
Replicate 2 enhancer activity (log2)
Replicate 1 enhancer activity (log2)
g.
minPs
Housekeeping
pGAPDH
pMYC
Developmental
pAPOBEC3F
eaBC
luc2
pA
ORFs
paBC
egfp
pA
Intrinsic minimal promoter activity
pGAPDH
pMYC
pAPOBEC3F
h.
All minPs
ρ = 0.75, P < 0.001
pGAPDH
pMYC
pAPOBEC3F
Enhancer activity (log2)
Promoter activity (log2)

**Supplementary Fig. 7: minP-TRE QUASARR-seq.**

**a-c**, Correlation of QUASARR-seq promoter activity measurements between replicates when paired with pGAPDH (a), pMYC (b), and pAPOBEC3F (c). **d-f**, Correlation of QUASARR-seq enhancer activity measurements between replicates when paired with pGAPDH (d), pMYC (e), and pAPOBEC3F (f). **g**, Top, QUASARR-seq pairing TREs with different minPs, including the promoters of a housekeeping gene; GAPDH, and a developmental gene; APOBEC3F, in addition to the promoter of MYC used thus far in this study. To distinguish between libraries, unique three-bp barcodes specific to each minP library were incorporated directly downstream of the paBCs. Bottom, Intrinsic promoter activities for the three minPs, calculated by taking their mean activities when paired with negative controls ORFs. Values shown are back-transformed from $\log_2$ scale. Error bars denote ±1 SEM. **h**, Correlation between element promoter and enhancer activities when paired with all minPs (Spearman's $\rho = 0.75$, $P$-value < 0.001).

a.

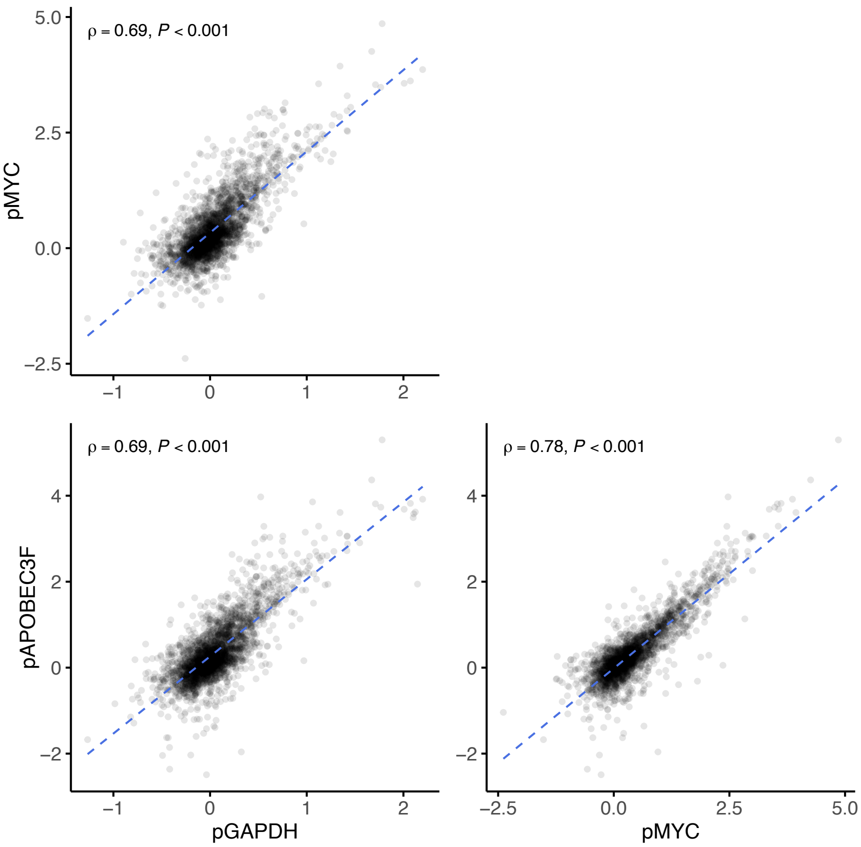

ρ = 0.69, P < 0.001
ρ = 0.69, P < 0.001
ρ = 0.78, P < 0.001
pMYC
pAPOBEC3F
pGAPDH
pMYC
5.0
2.5
0.0
−2.5
4
2
0
−2
−1
0
1
2
−2.5
0.0
2.5
5.0

**Supplementary Fig. 8: Enhancer-promoter compatibility is quantitative, not absolute.**
**a**, Pairwise correlations of QUASARR-seq enhancer activity measurements across minP libraries: pGAPDH vs. pMYC (Spearman's $\rho = 0.69$, *P*-value < 0.001; top left), pGAPDH vs. pAPOBEC3F ($\rho = 0.69$, *P*-value < 0.001; bottom left), and pMYC vs. pAPOBEC3F ($\rho = 0.78$, *P*-value < 0.001; bottom right).

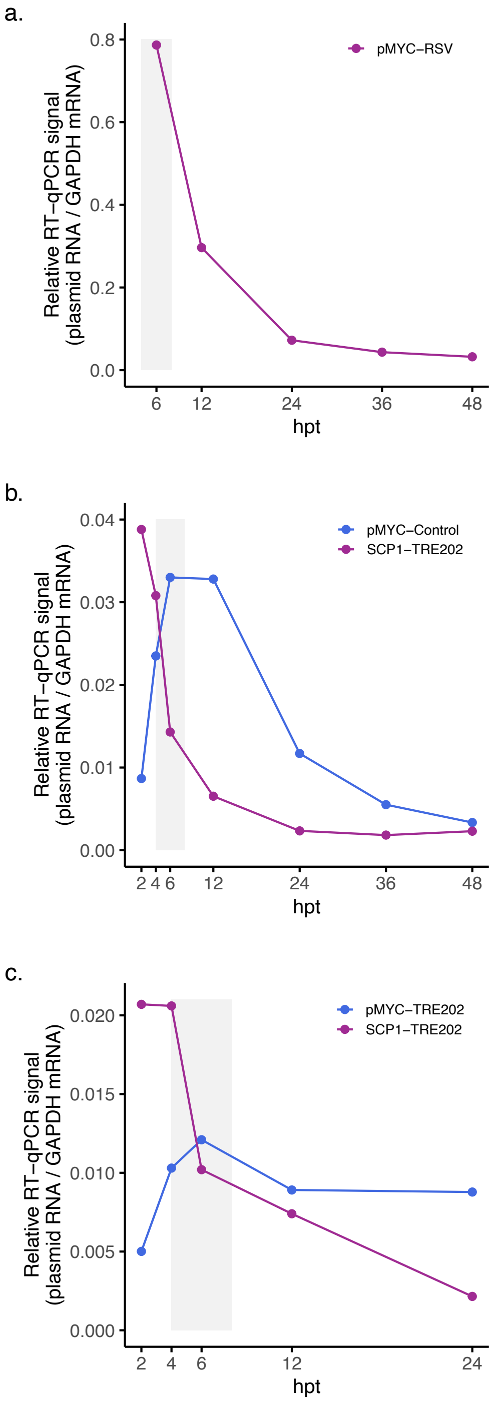

a.
Relative RT−qPCR signal
(plasmid RNA / GAPDH mRNA)
0.0
0.2
0.4
0.6
0.8
pMYC−RSV
6
12
24
36
48
hpt
b.
Relative RT−qPCR signal
(plasmid RNA / GAPDH mRNA)
0.00
0.01
0.02
0.03
0.04
pMYC−Control
SCP1−TRE202
2
4
6
12
24
36
48
hpt
c.
Relative RT−qPCR signal
(plasmid RNA / GAPDH mRNA)
0.000
0.005
0.010
0.015
0.020
pMYC−TRE202
SCP1−TRE202
2
4
6
12
24
hpt

**Supplementary Fig. 9: Optimal harvest time experiments.**

Relative RT-qPCR signal of plasmid-derived RNA normalized to GAPDH mRNA in K-562 transfected with **a**, pDEST-hSTARR-luc-pMYC-RSV, **b**, pDEST-hSTARR-luc-pMYC-Control1K or pDEST-hSTARR-luc-TRE202, and **c**, pDEST-hSTARR-luc-TRE202 or pDEST-hSTARR-luc-pMYC-TRE202 at varying hours post transfection (hpt). Points represent mean of technical triplicates per time point.
